# Specimen, Biological Structure, and Spatial Ontologies in Support of a Human Reference Atlas

**DOI:** 10.1101/2022.09.08.507220

**Authors:** Bruce W. Herr, Josef Hardi, Ellen M. Quardokus, Andreas Bueckle, Lu Chen, Fusheng Wang, Anita R. Caron, David Osumi-Sutherland, Mark A. Musen, Katy Börner

**Author notes:** Corresponding authors Katy Börner, Andreas Bueckle.

## Abstract

The Human Reference Atlas (HRA) is defined as a comprehensive, three-dimensional (3D) atlas of all the cells in the healthy human body. It is compiled by an international team of experts that develop standard terminologies linked to 3D reference objects describing anatomical structures. The third HRA release (v1.2) covers spatial reference data and ontology annotations for 26 organs. Experts access the HRA annotations via spreadsheets and view reference models in 3D editing tools. This paper introduces the Common Coordinate Framework Ontology (CCFO) v2.0.1 that interlinks specimen, biological structure, and spatial data together with the CCF API which makes the HRA programmatically accessible and interoperable with Linked Open Data (LOD). We detail how real-world user needs and experimental data guide CCFO design and implementation, present CCFO classes and properties together with examples of their usage, and report on technical validation performed. The CCFO graph database and API are used in the HuBMAP portal, Virtual Reality Organ Gallery, and other applications that support data queries across multiple, heterogeneous sources.

## Background & Summary

The Human BioMolecular Atlas Program (HuBMAP)^1^ is developing a human reference atlas (HRA) of the trillions of cells in the human body in close collaboration with 16 other consortia^2^. The HRA aims to serve the needs of biomedical researchers, practitioners, teachers and learners. It attempts to capture healthy human diversity (e.g., the location and type of cells and associated biomolecular data), and it will be useful to biomedical experts for many purposes. One major use case is allowing researchers to assess changes in the numbers and types of cells due to aging and disease.

The third HRA (v1.2) was released in June 2022, and it includes 26 organ-specific Anatomical Structures, Cell Types plus Biomarkers (ASCT+B) tables and 53 associated 3D reference organ models, among other features. On June 23, 2022, the HRA had been successfully used across four consortia to register 350 tissue blocks. On September 1, 2022, the count had increased to 364 tissue blocks and these tissue blocks link to 5,000 biomolecular datasets derived from these tissue samples.

### Human Reference Atlas

To construct the HRA, it is necessary to standardize experimental specimen metadata and to have a common coordinate framework (CCF)^3^ that provides a biological structure vocabulary linked to 3D spatial data for every organ, tissue, and cell in the human body^4^. To construct the CCF for the HRA, we assume that (1) anatomical structures are composed of cells that have different types (i.e., cell types are located in anatomical structures), combinations of biomarkers characterize different cell types, and experts are interested in querying for those entities (i.e., we need standardized terminology to query for anatomical structures, cell types, and biomarkers). We call this biological structure information. (2) The 3D space in which anatomical structures, cells of different types, and biomarkers operate does matter. We call this spatial information. (3) Human sex, age, and ethnicity impacts the structure and function of anatomical structures, cells, and biomarkers. We call this specimen information and use it to capture human diversity and its impact on body functions.

### Biological structure information

Anatomical structures, cell types, and biomarkers (ASCT+B) tables record the biomarker sets that characterize cell types within the nested anatomical structures that compose an organ. The ASCT+B tables are authored by human experts using templated Google Sheets—see standard operating procedure (SOP) entitled “SOP: Authoring ASCT+B Tables”^5^. The biomarkers, cell types, and anatomical structures are mapped to existing ontologies if they are available (e.g., to Uberon^6,7^). An exemplary ASCT+B table with an anatomical structure partonomy (in red on left) linked both to cell types (in blue in middle) and biomarkers (in green on right) is shown in **Fig. 1a**. All existing ASCT+B tables can be interactively explored in the ASCT+B Reporter^8^.

**Figure 1.**
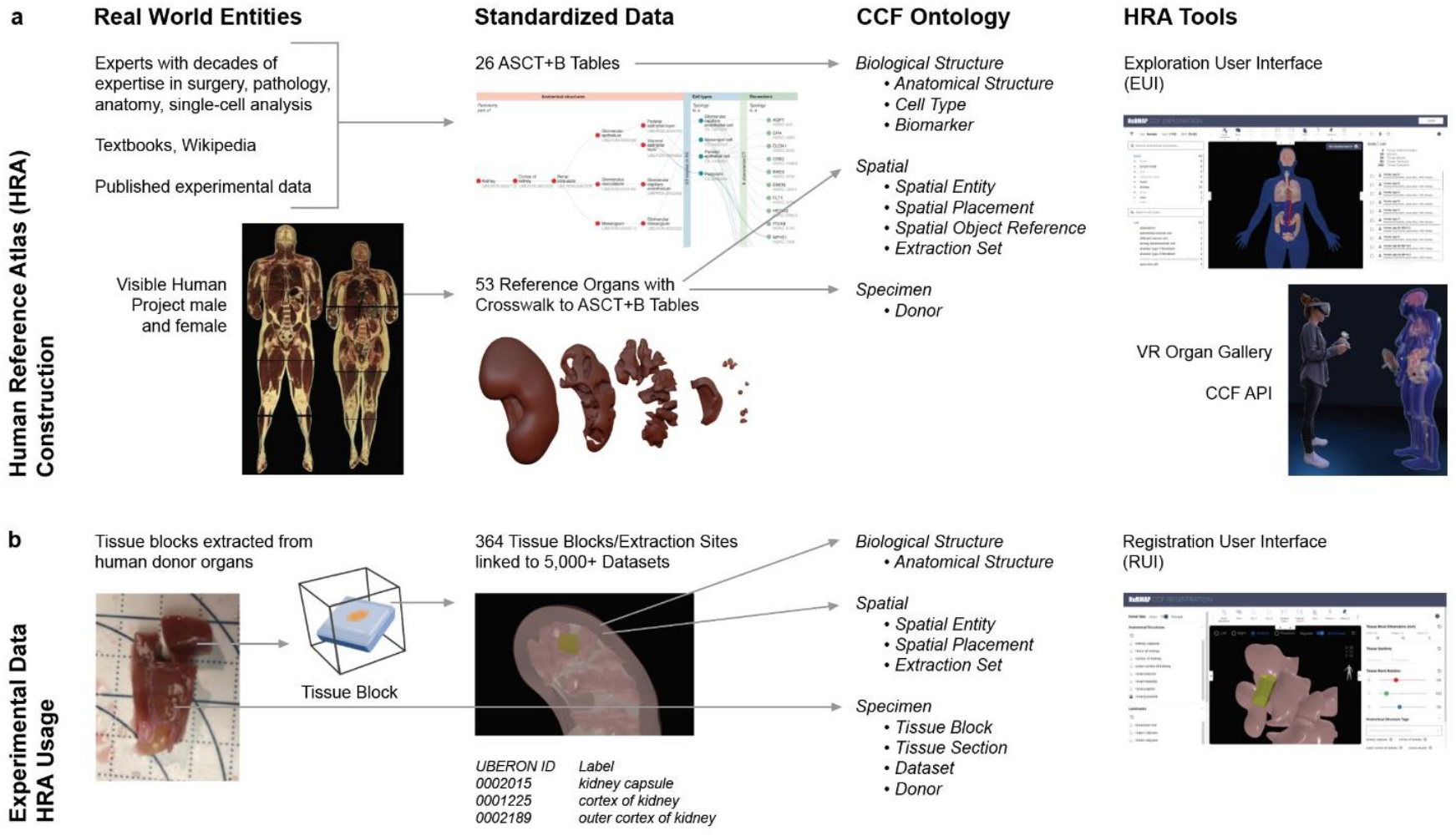
From real-world entities, to standardized data, to ontology. **a**. Human Reference Atlas construction takes real-world data and represents it in standardized data structures that are defined by the interlinked Biological Structure, Spatial, and Specimen ontologies. Anatomical structures in the ASCT+B tables are linked via *part_of* relationships resulting in a partonomy; they are crosswalked to 3D reference organs in support of spatial queries and exploration using different tools shown on the right. **b**. HRA usage includes search for specific cell types or biomarker expression values across organs in a 3D reference space. Users can also upload new experimental data (e.g., a new tissue block from a human donor specimen); the tissue is registered into the HRA by spatially mapping it to a 3D reference organ. If the Registration User Interface (RUI) shown on left is used, anatomical structure tags (see UBERON ID and Label) are automatically assigned based on collision events; spatial search becomes possible in the Exploration User Interface (EUI); and cell types or biomarkers associated with the colliding anatomical structures can be retrieved via ASCT+B tables.

### Spatial information

The 3D Reference Object Library captures the shape, size, location, and rotation of major anatomical structures. They are authored by medical illustrators following the “SOP: 3D Reference Object Approval”^9^. For each organ, there exists a digital representation of the so-called scene graph that represents all 3D objects (i.e., anatomical structures) and inter-object relationships (e.g., what object is inside or next to which other object). There exists a crosswalk table that associates 3D object names with the anatomical structure terms in the ASCT+B tables that link to terminology in existing ontologies whenever possible. **Fig. 1a** depicts how Visible Human Project data (VHP)^10^ (a high-quality, whole-body dataset published by the National Library of Medicine) is used to create reference organs (different anatomical structures of a kidney are shown) and how resulting data is uniformly represented using the spatial and specimen ontology. All reference organs can be interactively explored in the Exploration User Interface (EUI)^11^ and the Virtual Reality Organ Gallery^12^, both of which are detailed in the Applications section. The spatial size, shape, location, and rotation of experimental tissue data is captured relative to the organs in the 3D Reference Object Library. Uniform tissue registration is facilitated by the Registration User Interface (RUI) (**Fig. 2b, right**).

**Figure 2.**
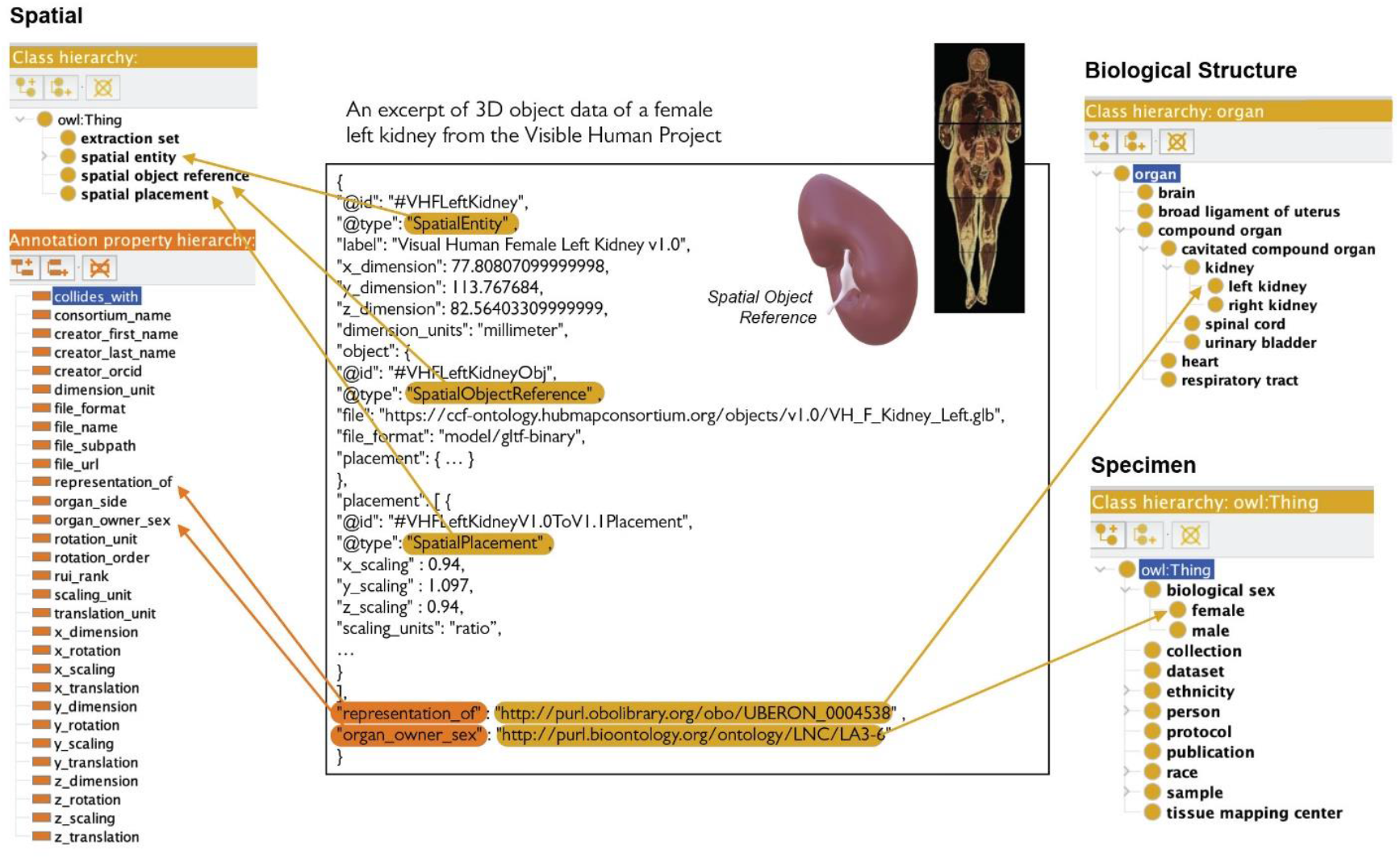
HRA Reference Organ Description. The Visible Human Project female left kidney (#VHFLeftKidney) reference organ is represented as a *Spatial Entity* class of type *Spatial Object Reference*, and with a *Spatial Placement*. Properties such as *representation_of* with PURL link to Uberon:0004538 (left kidney) refer to Biological Structure data, whereas properties such as *organ_owner_sex* with PURL link to LNC:LA3-6 (female) refer to Specimen data.

### Specimen information

Data capturing the sex, age, and other information on donors that provided tissue data used in the construction of the HRA are tracked. For the HRA (v1.2), data from the Visible Human Project was used in the construction of the 3D reference organs—the male was white, 180.3 cm (71 inch) tall, 90 kg (199 pound) and was 38 years old; the female was white, 171.2 cm (67.4 inch) tall, obese (weight unknown), and 59 years old.

### Provenance

We track scholarly publications and experimental studies that provide evidence for entities in the HRA (e.g., for what cell types have what gene expressions in which tissues). In addition to data provenance, we capture data processing provenance so the HRA can be recomputed, as needed, when new data and algorithms become available.

### HRA construction versus usage

Just as with geospatial maps, there is a difference between atlas creation (i.e., defining the reference space) and atlas usage (e.g., finding a place in the reference space or a pathway between two locations). **Fig. 1** provides an overview of HRA construction and usage by experts (e.g., anatomists, pathologists, surgeons, single cell researchers). We show real-world entities on the left, standardized data in the middle, and CCF ontologies and value-added services on the right. **Fig. 1a** shows HRA construction that includes the creation, validation, and publishing of the CCFO and associated APIs and tools. It starts with the creation of ASCT+B tables by experts with decades of expertise in surgery, pathology, anatomy, single-cell analysis, consulting textbooks, Wikipedia, and published experimental data. In parallel, professional medical illustrators create 3D spatial reference organs in support of tissue registration (e.g., the resulting reference organs have anatomical structures that serve as landmarks when registering virtual tissue blocks and can be used to assign semantic tags). Last but not least, crosswalks are compiled that link all 3D reference objects to anatomical structures in the ASCT+B tables (see details in Börner et al., 2021^4^). **Fig. 1b** details HRA usage—e.g., the mapping of new experimental tissue data into the HRA so it can be compared across tissue donors, laboratories, assay types or by using the HRA to query for cell types and the genes they express in different locations of the 3D human body. We show how tissue blocks are cut from human organs (and possibly further cut into tissue sections). The tissue blocks are then spatially registered into 3D reference organs using the Registration User Interface^13^, shown on right (and discussed in the Applications section), that assigns anatomical structure annotations (Uberon IDs and labels) via 3D object collision detection (see papers^14,15^ and the Applications section);

Assigned anatomical structures and data in the ASCT+B tables are then used to predict cell types and gene expressions for a tissue block. HRA-predicted data can then be compared to experimental data and the HRA data is revised as needed to capture healthy human diversity. Note that only the highest quality experimental data is used for HRA construction; all CCF-registered data can be explored within the HRA.

### Common Coordinate Framework Ontology

The CCF Ontology (CCFO) detailed in this paper formalizes and interlinks major classes that describe biological structure, spatial, and specimen information captured in the HRA. It thus provides a unifying vocabulary for HRA construction and usage—making it possible to ingest external data sources; supporting uniform tissue sample registration that includes the spatial positioning and semantic annotations within the 3D reference organs; and supporting user-formulated cross-domain queries over tissue donor properties, anatomical structures, cell types, biomarkers, and 3D space. The CCFO makes it possible to publish HRA data in the Linked Open Data (LOD) cloud^16^ in a Findable, Accessible, Interoperable and Reusable (FAIR)^17^ manner; the CCF API supports programmatic access to HRA data. We provide templates via data specifications so that other consortia can easily publish their data in a HRA-compatible manner.

This paper presents user needs that guide HRA and CCFO development. We then detail CCFO modeling and implementation as well as validation.

## Methods

### HRA Needs That Guide CCF Ontology Modeling

As stated above, there is a major difference between (1) the construction of the HRA (i.e., developing a latitude/longitude-like system) and agreeing on a way to map new data onto it and (2) the usage of the HRA to run different semantic and spatial queries. It is critically important to understand the goals and user needs for both parts to guide CCF Ontology design and HRA tool development (see examples in the Applications section). At their core, HRA queries act as competency questions^18,19^ that inform the modeling of the CCF Ontology.

#### HRA construction

Both human expertise and the highest quality experimental data are used to construct the HRA. Some data exists only at the gross anatomy level (e.g., data covered in anatomy books or Visible Human Project data), while other data covers single-cell data for micro-anatomical structures (e.g., gene expression data for cells). All data—not only human expertise but also experimental data across different levels of resolution—must be registered into the very same HRA CCF. To register experimental data, hundreds of thousands of tissue blocks (currently ca. 5,000) collected from thousands of human donors (currently ca. 200) must be uniformly registered and annotated in terms of biological structure, spatial, and specimen data. HRA terms should link to existing ontologies whenever possible. If HRA terms do not exist in reference ontologies, then a GitHub issue is submitted to the respective ontology for eventual inclusion of these anatomical structures and cell types into existing ontologies and can be tracked^20,21^. Collision detection at the mesh level using the 3D Geometry-Based Tissue Block Annotation tool^14^ (see Applications section) needs to be run to identify what type and volume of anatomical structures a tissue block contains (this step is necessary as most tissue samples cover cells from multiple anatomical structures). Datasets for these tissue blocks (e.g., cell-type name annotations computed using the Azimuth^22^ tool or gene-expression values measured for cells in experiments) must be mapped to anatomical structures—so it becomes possible to show cell-type populations (i.e., the number of cells per type within an anatomical structure) inside of each 3D anatomical structure. For each HRA entity, all data and code needed to recompile it must be tracked to ensure reproducibility. A new HRA is computed over the course of several days using massive storage and computing resources every six months. Upon each successful recompilation, baseline HRA statistics such as entity and relationship counts, cell-type populations, or query-performance test results are computed and published.

#### HRA usage

Some users are interested to understand in which 3D spatial locations or in what anatomical structures cells of a certain type (e.g., immune cells) can be found or what biomarkers are available or common for a certain cell type (e.g., are there differences in gene expression values for a certain cell type across organs). Other users would like to compare their very own experimental data to the growing set of HRA compatible data (e.g., to understand how diseased cell-type populations or biomarker-expression values differ from a healthy reference). A third set of users is interested in transferring cell-type annotations from the HRA to a new experimental dataset (e.g., using the Azimuth^22^ tool that automates the analysis, interpretation of single-cell RNA-seq experiment data and transfers cell annotations from the reference map to new experimental data). If new data is uploaded, it needs to be registered and analyzed at scale; 3D exploration requires that 3D reference organ files are loaded efficiently and spatial and other queries are run fast using specimen, biological structure, and/or spatial metadata. For example, a user might like to see all tissue blocks for 50-year-old females or search for cell types in a specific spatial location that have high gene-expression values in experimental datasets. The interlinkage of spatial reference objects and ASCT+B tables makes it possible to associate cell-type and biomarker data from experimental data to 3D anatomical structure data and to compute how many cells of what type are commonly found in which anatomical structures and what their typical biomarker expression values are. As the number of HRA-registered datasets increases, prediction accuracy will increase. As the HRA is spatially explicit, the spatial location of cell-type populations can be visually presented in 3D. Initially, cells are placed randomly; as more data on cell-neighborhoods becomes available^23,24^, cells can be placed in spatially explicit patterns. Usage statistics must be captured to identify bottlenecks (e.g., long loading or computing times) and to improve application performance.

Note that the HRA is continuously evolving as new, higher-quality datasets become available for an increasing number of organs. HRA construction and optimization for different use cases will likely take decades. As with any atlas, the availability of the HRA will inspire qualitatively novel questions and queries that are hard to envision today.

### CCF Ontology Modeling and Design

The CCFO v2.0.1 focuses on the subset of the third HRA (v1.2) release that is currently used for tissue registration and exploration in the HuBMAP^25^ and GTEx^26^ data portals and in other tools and services discussed in the Applications section.

#### CCFO Modeling Assumptions and Goals

The CCFO is modeled to serve the needs of HRA construction and usage. It formally defines the three interlinked parts of the HRA via three interlinked ontologies called Biological Structure, Spatial, and Specimen Ontology (see **Fig. 1**). Here, we discuss CCFO modeling assumptions, goals, and desirable properties.

##### FAIR and interoperable

We want the HRA to be findable and interoperable with other high-quality data sources outside the HRA. Therefore, HRA terms should be linked, whenever possible, to existing, widely used ontologies. For example, HRA anatomical structures are linked to Uberon^6,7^ (and FMA^27–29^ as needed) and cell types are linked to the Cell Ontology^30,31^. These two ontologies are used extensively to annotate single-cell transcriptomics data^32,33^ and are cross integrated with each other and with other widely used ontologies (e.g., the Gene Ontology [GO]^34^, the Human Phenotype Ontology^35^, and the Mondo Disease Ontology^36^). HRA gene and protein biomarkers are linked to HUGO Gene Nomenclature Committee (HGNC)^37^ and crosswalks exist to Ensembl^38^ for genes and to UniProt^39^ for proteins.

##### Limitations of existing ontologies

Uberon and CL are overly complex for the needs of the HRA as they cover multiple species and developmental stages, and both ontologies lack some terms needed by the HRA. FMA can be hard to query due to inconsistent application of ontology design patterns, and it does not link to other ontologies. Existing reference ontologies cannot represent the spatial size, location, and rotation of 3D human reference organs and experimental data relative to the HRA and to each other; e.g., such as Uberon and FMA, have incorporated anatomical location descriptors using spatial properties (e.g., *anterior_to, posterior_to, left_of, right_of, opposite_to*, etc.), but these are often too general (e.g., in Uberon the location descriptors are for multiple species).

##### Use existing standards

CCFO defines a common vocabulary to annotate data in an HRA-compliant manner using several well-known biomedical ontologies, such as Uberon, FMA, CL, HGNC, and other semantic web vocabularies, such as the Dublin Core™ Metadata Initiative (DCTERMS)^40^ and RDFS^41^. The CCFO uses some Dublin Core properties for providing provenance information, such as the data publisher, creator, and creation date. The CCFO uses the Open Biological and Biomedical Ontology (OBO) for properties such as *Definitions* for anatomical structures, cell types, and biomarkers. CCFO is published using the standard Semantic Web technology OWL which is built on the Resource Description Framework (RDF) data format that was designed for interoperability and supporting inference capabilities that can be used for consistency checking in the future.

##### Validation

The anatomical structures and cell types in the ASCT+B tables are validated against reference ontologies on a weekly basis (see Technical Validation section). Validation results are used to improve the ASCT+B tables and to issue ontology change requests to slowly and steadily modify existing reference ontologies (e.g., Uberon and CL) so they accurately represent healthy human adult data. When completed, the CCFO will be a proper, HRA-labeled subset (using the *human_reference_atlas* subset tag) of major reference ontologies that are widely used by a diverse range of bioinformatics applications.

##### HRA will evolve over time

As new experimental datasets and analysis results become available, the HRA data structures and the CCFO will need to be updated to capture new entity types and linkages and to serve qualitatively new user needs.

#### CCFO Classes and Properties

The CCF ontology published as CCF.OWL v2.0.1^42^ comprises three interlinked ontologies that capture Biological Structure, Spatial, Specimen data (see **Fig. 1**). The names of the three ontologies, their classes, properties, and definitions can be found in **Table 1**.

**Table 1.**
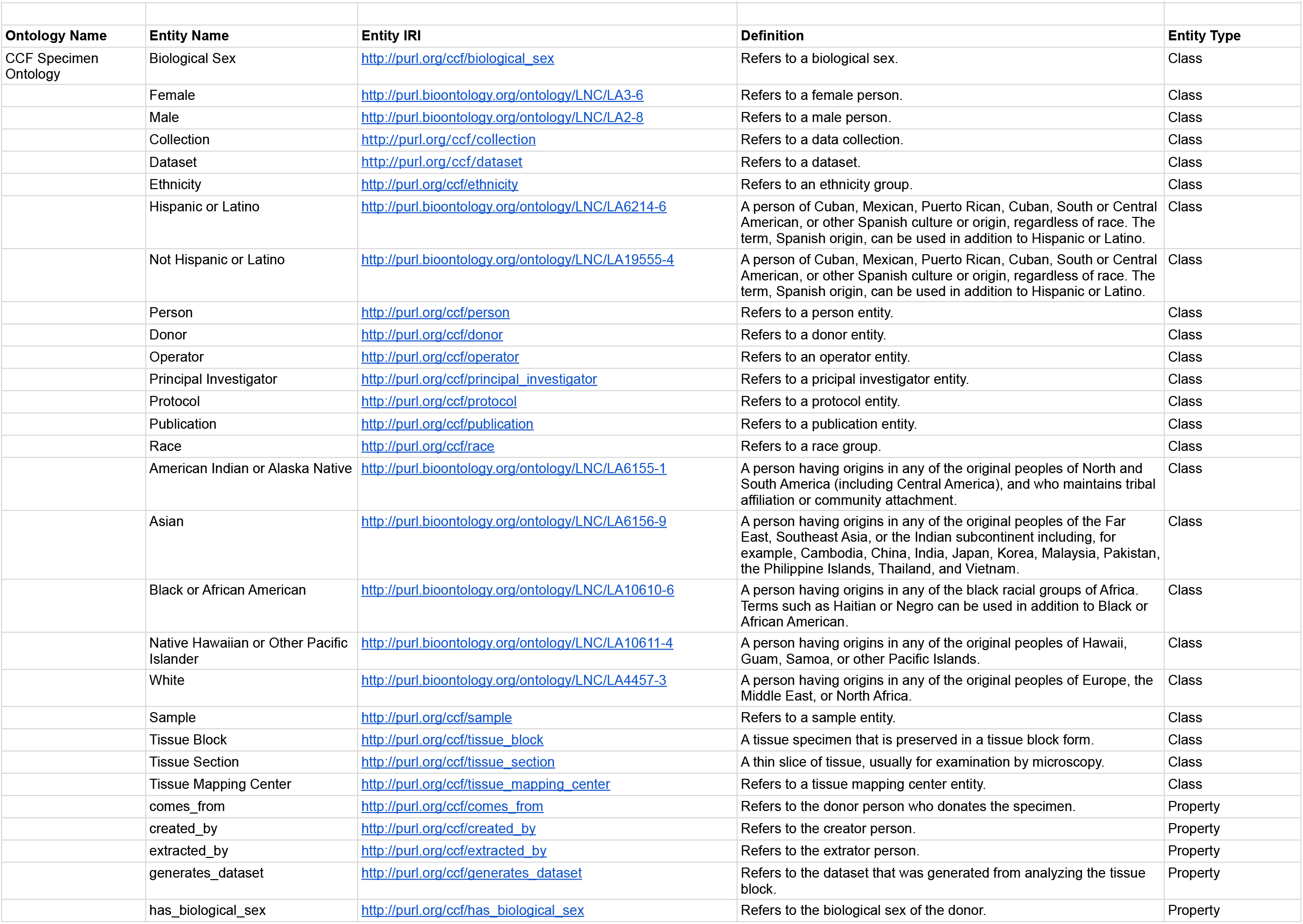
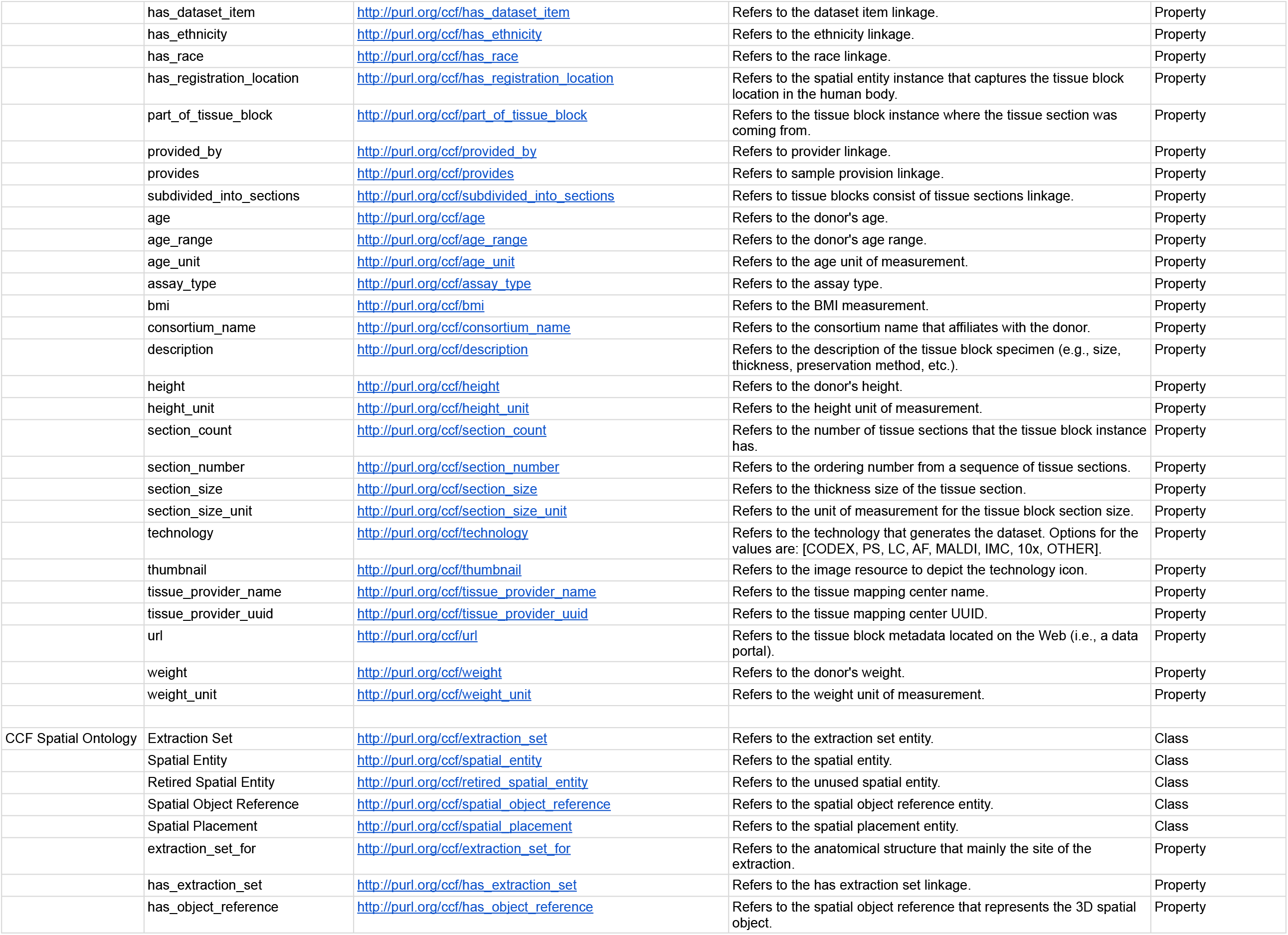
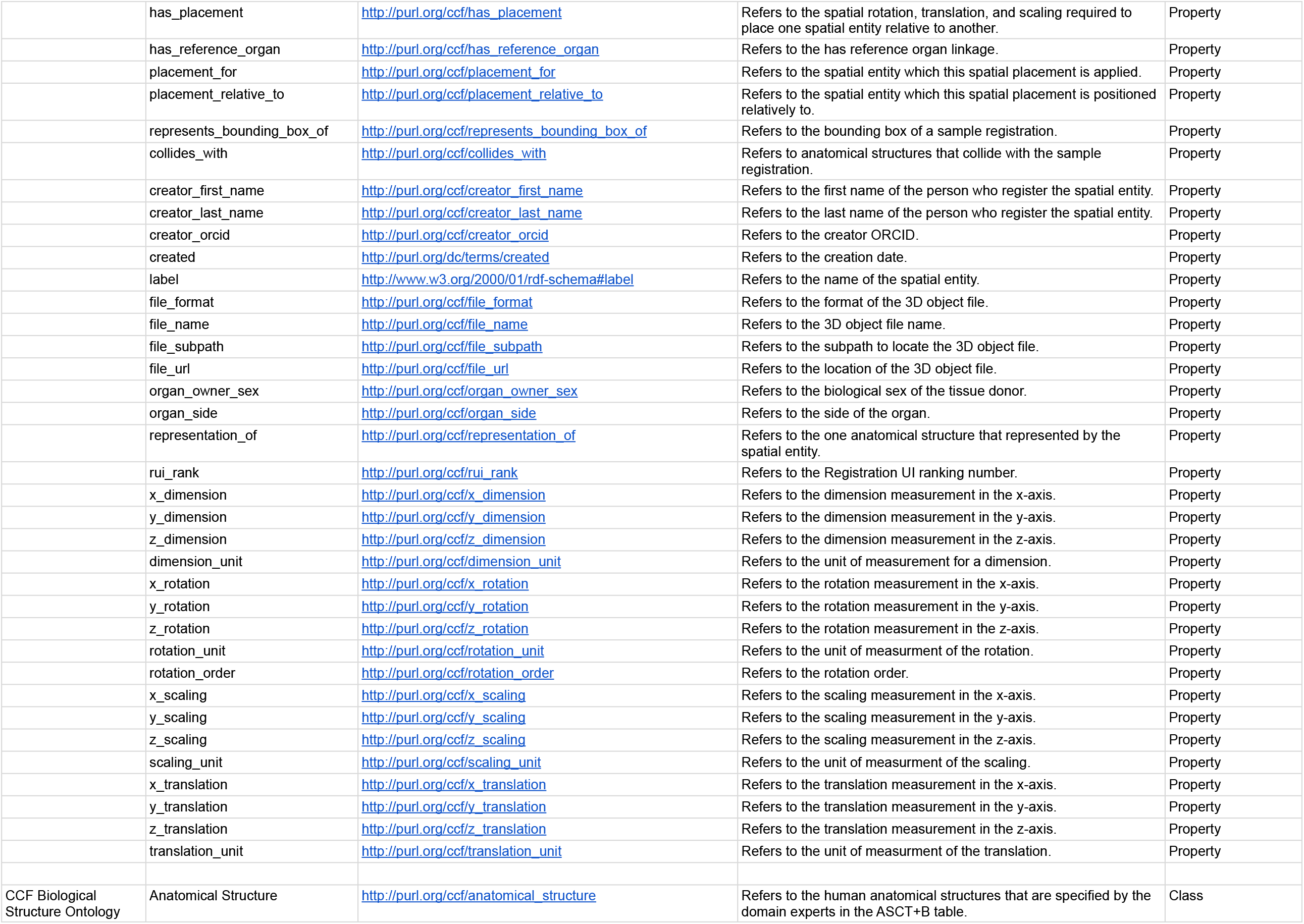
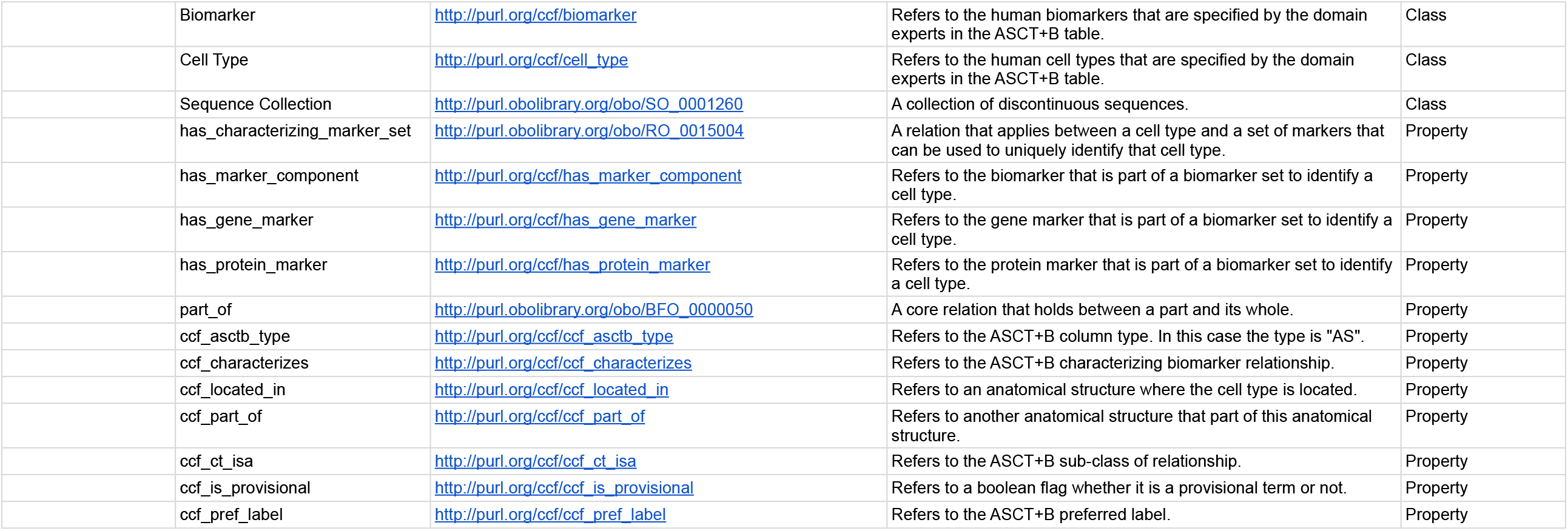
CCF Specimen, Biological Structure, and Spatial Ontology. For each ontology, we provide all class names, property names, entity Internationalized Resource Identifier (IRI), definition when available, and entity type.

| Ontology Name | Entity Name | Entity IRI | Definition | Entity Type |
| --- | --- | --- | --- | --- |
| CCF Specimen Ontology | Biological Sex | http://purl.org/ccf/biological_sex | Refers to a biological sex. | Class |
|  | Female | http://purl.bioontology.org/ontology/LNC/LA3-6 | Refers to a female person. | Class |
|  | Male | http://purl.bioontology.org/ontology/LNC/LA2-8 | Refers to a male person. | Class |
|  | Collection | http://purl.org/ccf/collection | Refers to a data collection. | Class |
|  | Dataset | http://purl.org/ccf/dataset | Refers to a dataset. | Class |
|  | Ethnicity | http://purl.org/ccf/ethnicity | Refers to an ethnicity group. | Class |
|  | Hispanic or Latino | http://purl.bioontology.org/ontology/LNC/LA6214-6 | A person of Cuban, Mexican, Puerto Rican, Cuban, South or Central American, or other Spanish culture or origin, regardless of race. The term, Spanish origin, can be used in addition to Hispanic or Latino. | Class |
|  | Not Hispanic or Latino | http://purl.bioontology.org/ontology/LNC/LA19555-4 | A person of Cuban, Mexican, Puerto Rican, Cuban, South or Central American, or other Spanish culture or origin, regardless of race. The term, Spanish origin, can be used in addition to Hispanic or Latino. | Class |
|  | Person | http://purl.org/ccf/person | Refers to a person entity. | Class |
|  | Donor | http://purl.org/ccf/donor | Refers to a donor entity. | Class |
|  | Operator | http://purl.org/ccf/operator | Refers to an operator entity. | Class |
|  | Principal Investigator | http://purl.org/ccf/principal_investigator | Refers to a principal investigator entity. | Class |
|  | Protocol | http://purl.org/ccf/protocol | Refers to a protocol entity. | Class |
|  | Publication | http://purl.org/ccf/publication | Refers to a publication entity. | Class |
|  | Race | http://purl.org/ccf/race | Refers to a race group. | Class |
|  | American Indian or Alaska Native | http://purl.bioontology.org/ontology/LNC/LA6155-1 | A person having origins in any of the original peoples of North and South America (including Central America), and who maintains tribal affiliation or community attachment. | Class |
|  | Asian | http://purl.bioontology.org/ontology/LNC/LA6156-9 | A person having origins in any of the original peoples of the Far East, Southeast Asia, or the Indian subcontinent including, for example, Cambodia, China, India, Japan, Korea, Malaysia, Pakistan, the Philippine Islands, Thailand, and Vietnam. | Class |
|  | Black or African American | http://purl.bioontology.org/ontology/LNC/LA10610-6 | A person having origins in any of the black racial groups of Africa. Terms such as Haitian or Negro can be used in addition to Black or African American. | Class |
|  | Native Hawaiian or Other Pacific Islander | http://purl.bioontology.org/ontology/LNC/LA10611-4 | A person having origins in any of the original peoples of Hawaii, Guam, Samoa, or other Pacific Islands. | Class |
|  | White | http://purl.bioontology.org/ontology/LNC/LA4457-3 | A person having origins in any of the original peoples of Europe, the Middle East, or North Africa. | Class |
|  | Sample | http://purl.org/ccf/sample | Refers to a sample entity. | Class |
|  | Tissue Block | http://purl.org/ccf/tissue_block | A tissue specimen that is preserved in a tissue block form. | Class |
|  | Tissue Section | http://purl.org/ccf/tissue_section | A thin slice of tissue, usually for examination by microscopy. | Class |
|  | Tissue Mapping Center | http://purl.org/ccf/tissue_mapping_center | Refers to a tissue mapping center entity. | Class |
|  | comes_from | http://purl.org/ccf/comes_from | Refers to the donor person who donates the specimen. | Property |
|  | created_by | http://purl.org/ccf/created_by | Refers to the creator person. | Property |
|  | extracted_by | http://purl.org/ccf/extracted_by | Refers to the extractor person. | Property |
|  | generates_dataset | http://purl.org/ccf/generates_dataset | Refers to the dataset that was generated from analyzing the tissue block. | Property |
|  | has_biological_sex | http://purl.org/ccf/has_biological_sex | Refers to the biological sex of the donor. | Property |
|  | has_dataset_item | http://purl.org/ccf/has_dataset_item | Refers to the dataset item linkage. | Property |
|  | has_ethnicity | http://purl.org/ccf/has_ethnicity | Refers to the ethnicity linkage. | Property |
|  | has_race | http://purl.org/ccf/has_race | Refers to the race linkage. | Property |
|  | has_registration_location | http://purl.org/ccf/has_registration_location | Refers to the spatial entity instance that captures the tissue block location in the human body. | Property |
|  | part_of_tissue_block | http://purl.org/ccf/part_of_tissue_block | Refers to the tissue block instance where the tissue section was coming from. | Property |
|  | provided_by | http://purl.org/ccf/provided_by | Refers to provider linkage. | Property |
|  | provides | http://purl.org/ccf/provides | Refers to sample provision linkage. | Property |
|  | subdivided_into_sections | http://purl.org/ccf/subdivided_into_sections | Refers to tissue blocks consist of tissue sections linkage. | Property |
|  | age | http://purl.org/ccf/age | Refers to the donor's age. | Property |
|  | age_range | http://purl.org/ccf/age_range | Refers to the donor's age range. | Property |
|  | age_unit | http://purl.org/ccf/age_unit | Refers to the age unit of measurement. | Property |
|  | assay_type | http://purl.org/ccf/assay_type | Refers to the assay type. | Property |
|  | bmi | http://purl.org/ccf/bmi | Refers to the BMI measurement. | Property |
|  | consortium_name | http://purl.org/ccf/consortium_name | Refers to the consortium name that affiliates with the donor. | Property |
|  | description | http://purl.org/ccf/description | Refers to the description of the tissue block specimen (e.g., size, thickness, preservation method, etc.). | Property |
|  | height | http://purl.org/ccf/height | Refers to the donor's height. | Property |
|  | height_unit | http://purl.org/ccf/height_unit | Refers to the height unit of measurement. | Property |
|  | section_count | http://purl.org/ccf/section_count | Refers to the number of tissue sections that the tissue block instance has. | Property |
|  | section_number | http://purl.org/ccf/section_number | Refers to the ordering number from a sequence of tissue sections. | Property |
|  | section_size | http://purl.org/ccf/section_size | Refers to the thickness size of the tissue section. | Property |
|  | section_size_unit | http://purl.org/ccf/section_size_unit | Refers to the unit of measurement for the tissue block section size. | Property |
|  | technology | http://purl.org/ccf/technology | Refers to the technology that generates the dataset. Options for the values are: [CODEX, PS, LC, AF, MALDI, IMC, 10x, OTHER]. | Property |
|  | thumbnail | http://purl.org/ccf/thumbnail | Refers to the image resource to depict the technology icon. | Property |
|  | tissue_provider_name | http://purl.org/ccf/tissue_provider_name | Refers to the tissue mapping center name. | Property |
|  | tissue_provider_uuid | http://purl.org/ccf/tissue_provider_uuid | Refers to the tissue mapping center UUID. | Property |
|  | url | http://purl.org/ccf/url | Refers to the tissue block metadata located on the Web (i.e., a data portal). | Property |
|  | weight | http://purl.org/ccf/weight | Refers to the donor's weight. | Property |
|  | weight_unit | http://purl.org/ccf/weight_unit | Refers to the weight unit of measurement. | Property |
| CCF Spatial Ontology | Extraction Set | http://purl.org/ccf/extraction_set | Refers to the extraction set entity. | Class |
|  | Spatial Entity | http://purl.org/ccf/spatial_entity | Refers to the spatial entity. | Class |
|  | Retired Spatial Entity | http://purl.org/ccf/retired_spatial_entity | Refers to the unused spatial entity. | Class |
|  | Spatial Object Reference | http://purl.org/ccf/spatial_object_reference | Refers to the spatial object reference entity. | Class |
|  | Spatial Placement | http://purl.org/ccf/spatial_placement | Refers to the spatial placement entity. | Class |
|  | extraction_set_for | http://purl.org/ccf/extraction_set_for | Refers to the anatomical structure that mainly the site of the extraction. | Property |
|  | has_extraction_set | http://purl.org/ccf/has_extraction_set | Refers to the has extraction set linkage. | Property |
|  | has_object_reference | http://purl.org/ccf/has_object_reference | Refers to the spatial object reference that represents the 3D spatial object. | Property |
|  | has_placement | http://purl.org/ccf/has_placement | Refers to the spatial rotation, translation, and scaling required to place one spatial entity relative to another. | Property |
|  | has_reference_organ | http://purl.org/ccf/has_reference_organ | Refers to the has reference organ linkage. | Property |
|  | placement_for | http://purl.org/ccf/placement_for | Refers to the spatial entity which this spatial placement is applied. | Property |
|  | placement_relative_to | http://purl.org/ccf/placement_relative_to | Refers to the spatial entity which this spatial placement is positioned relatively to. | Property |
|  | represents_bounding_box_of | http://purl.org/ccf/represents_bounding_box_of | Refers to the bounding box of a sample registration. | Property |
|  | collides_with | http://purl.org/ccf/collides_with | Refers to anatomical structures that collide with the sample registration. | Property |
|  | creator_first_name | http://purl.org/ccf/creator_first_name | Refers to the first name of the person who register the spatial entity. | Property |
|  | creator_last_name | http://purl.org/ccf/creator_last_name | Refers to the last name of the person who register the spatial entity. | Property |
|  | creator_orcid | http://purl.org/ccf/creator_orcid | Refers to the creator ORCID. | Property |
|  | created | http://purl.org/dc/terms/created | Refers to the creation date. | Property |
|  | label | http://www.w3.org/2000/01/rdf-schema#label | Refers to the name of the spatial entity. | Property |
|  | file_format | http://purl.org/ccf/file_format | Refers to the format of the 3D object file. | Property |
|  | file_name | http://purl.org/ccf/file_name | Refers to the 3D object file name. | Property |
|  | file_subpath | http://purl.org/ccf/file_subpath | Refers to the subpath to locate the 3D object file. | Property |
|  | file_url | http://purl.org/ccf/file_url | Refers to the location of the 3D object file. | Property |
|  | organ_owner_sex | http://purl.org/ccf/organ_owner_sex | Refers to the biological sex of the tissue donor. | Property |
|  | organ_side | http://purl.org/ccf/organ_side | Refers to the side of the organ. | Property |
|  | representation_of | http://purl.org/ccf/representation_of | Refers to the one anatomical structure that represented by the spatial entity. | Property |
|  | rui_rank | http://purl.org/ccf/rui_rank | Refers to the Registration UI ranking number. | Property |
|  | x_dimension | http://purl.org/ccf/x_dimension | Refers to the dimension measurement in the x-axis. | Property |
|  | y_dimension | http://purl.org/ccf/y_dimension | Refers to the dimension measurement in the y-axis. | Property |
|  | z_dimension | http://purl.org/ccf/z_dimension | Refers to the dimension measurement in the z-axis. | Property |
|  | dimension_unit | http://purl.org/ccf/dimension_unit | Refers to the unit of measurement for a dimension. | Property |
|  | x_rotation | http://purl.org/ccf/x_rotation | Refers to the rotation measurement in the x-axis. | Property |
|  | y_rotation | http://purl.org/ccf/y_rotation | Refers to the rotation measurement in the y-axis. | Property |
|  | z_rotation | http://purl.org/ccf/z_rotation | Refers to the rotation measurement in the z-axis. | Property |
|  | rotation_unit | http://purl.org/ccf/rotation_unit | Refers to the unit of measurment of the rotation. | Property |
|  | rotation_order | http://purl.org/ccf/rotation_order | Refers to the rotation order. | Property |
|  | x_scaling | http://purl.org/ccf/x_scaling | Refers to the scaling measurement in the x-axis. | Property |
|  | y_scaling | http://purl.org/ccf/y_scaling | Refers to the scaling measurement in the y-axis. | Property |
|  | z_scaling | http://purl.org/ccf/z_scaling | Refers to the scaling measurement in the z-axis. | Property |
|  | scaling_unit | http://purl.org/ccf/scaling_unit | Refers to the unit of measurment of the scaling. | Property |
|  | x_translation | http://purl.org/ccf/x_translation | Refers to the translation measurement in the x-axis. | Property |
|  | y_translation | http://purl.org/ccf/y_translation | Refers to the translation measurement in the y-axis. | Property |
|  | z_translation | http://purl.org/ccf/z_translation | Refers to the translation measurement in the z-axis. | Property |
|  | translation_unit | http://purl.org/ccf/translation_unit | Refers to the unit of measurment of the translation. | Property |
| CCF Biological Structure Ontology | Anatomical Structure | http://purl.org/ccf/anatomical_structure | Refers to the human anatomical structures that are specified by the domain experts in the ASCT+B table. | Class |

|  |  |  |  |
| --- | --- | --- | --- |
| Biomarker | http://purl.org/ccf/biomarker | Refers to the human biomarkers that are specified by the domain experts in the ASCT+B table. | Class |
| Cell Type | http://purl.org/ccf/cell_type | Refers to the human cell types that are specified by the domain experts in the ASCT+B table. | Class |
| Sequence Collection | http://purl.obolibrary.org/obo/SO_0001260 | A collection of discontinuous sequences. | Class |
| has_characterizing_marker_set | http://purl.obolibrary.org/obo/RO_0015004 | A relation that applies between a cell type and a set of markers that can be used to uniquely identify that cell type. | Property |
| has_marker_component | http://purl.org/ccf/has_marker_component | Refers to the biomarker that is part of a biomarker set to identify a cell type. | Property |
| has_gene_marker | http://purl.org/ccf/has_gene_marker | Refers to the gene marker that is part of a biomarker set to identify a cell type. | Property |
| has_protein_marker | http://purl.org/ccf/has_protein_marker | Refers to the protein marker that is part of a biomarker set to identify a cell type. | Property |
| part_of | http://purl.obolibrary.org/obo/BFO_0000050 | A core relation that holds between a part and its whole. | Property |
| ccf_asctb_type | http://purl.org/ccf/ccf_asctb_type | Refers to the ASCT+B column type. In this case the type is "AS". | Property |
| ccf_characterizes | http://purl.org/ccf/ccf_characterizes | Refers to the ASCT+B characterizing biomarker relationship. | Property |
| ccf_located_in | http://purl.org/ccf/ccf_located_in | Refers to an anatomical structure where the cell type is located. | Property |
| ccf_part_of | http://purl.org/ccf/ccf_part_of | Refers to another anatomical structure that part of this anatomical structure. | Property |
| ccf_ct_isa | http://purl.org/ccf/ccf_ct_isa | Refers to the ASCT+B sub-class of relationship. | Property |
| ccf_is_provisional | http://purl.org/ccf/ccf_is_provisional | Refers to a boolean flag whether it is a provisional term or not. | Property |
| ccf_pref_label | http://purl.org/ccf/ccf_pref_label | Refers to the ASCT+B preferred label. | Property |

**The Biological Structure Ontology** has three main classes: *Anatomical Structure, Cell Type*, and *Biomarker* that are derived from the ASCT+B tables for HRA construction. During HRA usage, *Anatomical Structure* data is captured via collision events and from running different assay types. In the ASCT+B tables, *Anatomical Structure* terms are linked to Uberon (supplementing with FMA^27–29^ as needed). *Cell Type* terms link mainly to CL^30,31^ (supplementing with Provisional Cell Ontology (PCL) for brain^43^ and Human Lung Maturation (LungMAP Human Anatomy (LMHA)^44^ for lung. Note that Uberon and CL are already widely used by the single-cell research community making them an obvious choice. *Biomarker* terms are linked to HGNC^37^ standard pipeline developed by the brain data standards initiative^43^ and is used to link these to cell-type terms.

**The Spatial Ontology** has four main classes: *Spatial Entity, Spatial Placement, Spatial Object Reference*, and *Extraction Set*. HRA Construction uses all four classes, while HRA Usage uses the first two only. Terms for anatomical structures from the Biological Structure Ontology are linked to 3D reference data in a CCF in support of spatial registration, spatial search among others. Concretely, each object in a 3D organ is annotated with the appropriate anatomical structure term from the partonomy tree in the Biological Structure Ontology. Lastly, each 3D organ object instance in the ontology receives a set of spatial placement parameters that states its position, size, and orientation in relation to the HRA coordinate space.

**The Specimen Ontology** has four main classes: *Tissue Block, Tissue Section, Dataset, Donor*. Only donor is used for HRA construction; all four are used for HRA usage. For HRA construction, we keep track of the Visible Human Project specimen data. For HRA usage, we record *Tissue Block, Tissue Section, Dataset, Donor* info for each tissue block. In terms of *Donor* data that is collected by independent laboratories, we capture sex and age, among other properties.

#### CCFO Examples

This section presents examples of how the ASCT+B tables, 3D reference organs, landmarks, and experimental tissue block data are represented using CCFO classes and properties. We render ontology class names in mixed case *italics* (e.g., *Spatial Entity*, where the first letter of each compound word is capitalized), set ontology properties in lower-case *italics* with underscores between words (e.g., *part_of* or *has_object_reference*), and use bold italics for OWL Web Ontology Language reserved words (e.g., ***SubClassOf***). We use normal orthography to describe what a class or property represents.

##### HRA reference organs

An excerpt of how a reference organ is represented using the CCFO is shown in **Fig. 2**. The middle panel text details how the Visible Human Project female left kidney (#VHFLeftKidney) reference organ is represented as a *Spatial Entity*. This *Spatial Entity* has well-defined 3D spatial dimensions and a link to a GL Transmission Format Binary (GLB) object file of type *Spatial Object Reference* that captures the 3D shape of this anatomical structure.

The reference object is properly placed within the female reference body via a *Spatial Placement* that defines 3D scaling among others. Highlighted in orange are *representation_of* and *organ_owner_sex* that are encoded as Annotation Properties (in lower left) with persistent uniform resource locator (PURL) links to Uberon ID 0004538 for “left kidney” in the Biological Structure Ontology (top right) and to the Bioontology Logical Observation Identifier Names and Codes ID LA3-6 for “female” sex in the Specimen Ontology (in lower right).

##### Experimental tissue data

The CCFO is also used to represent experimental data. **Fig. 3** exemplifies how data for a *Sample* of type *Tissue Block* that was provided by a *Donor* which is a *Person* in the Specimen Ontology (in lower right). The *Tissue Block* inherits the *has_registration_location* from a *Spatial Entity* with property *has_placement* inside of the #VHFLeftKidney reference organ given in **Fig. 2**. 3D object collision events (commonly used in computer games to compute spatial relationships among moving objects) are used to compute which anatomical structures are inside of a tissue block (see Applications section for details).

**Figure 3.**
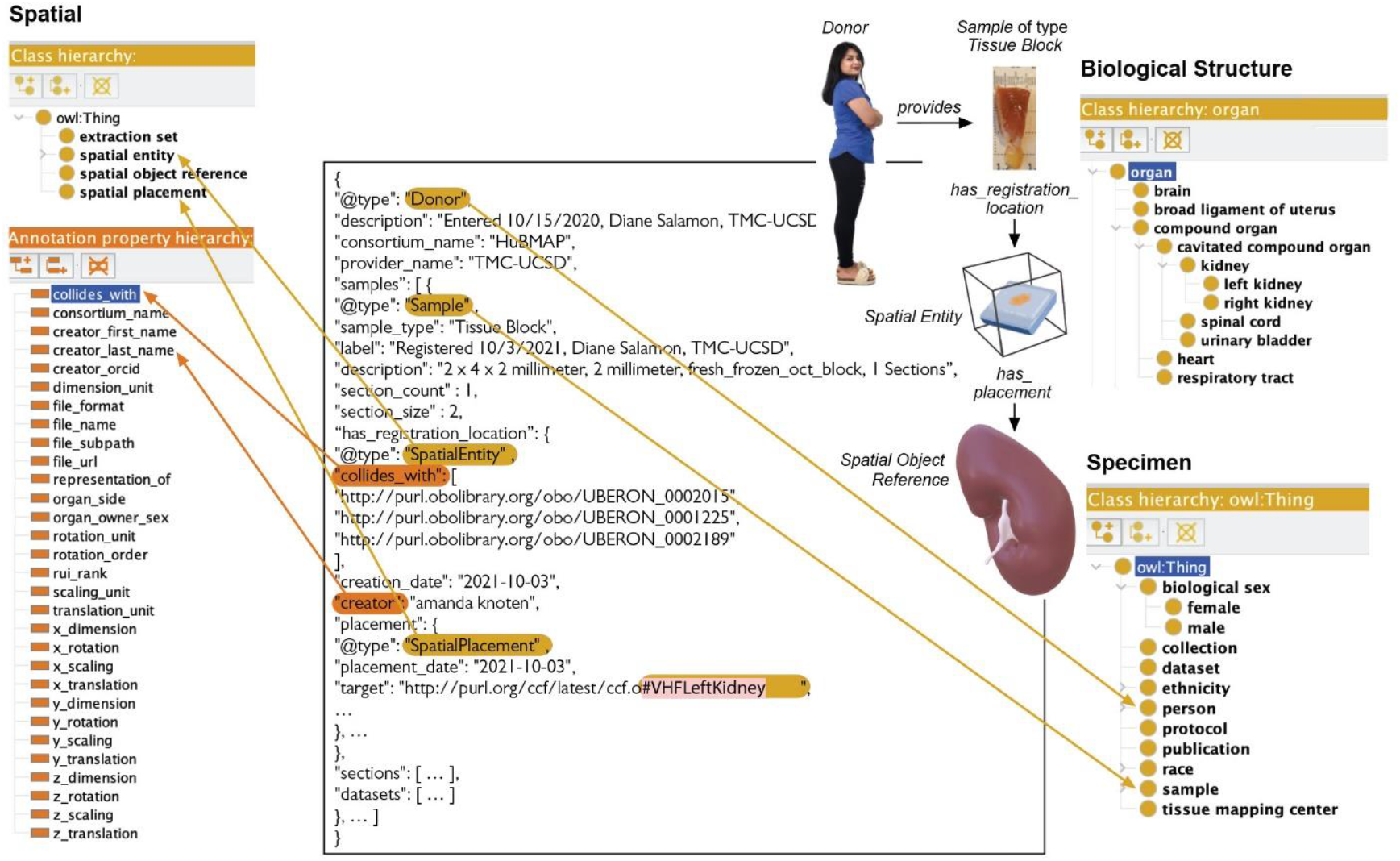
HRA Tissue Block Description. A *Donor* (which *is_a Person*) provides a *Sample* of type *Tissue Block* with properties such as *Biological Sex* and annotation properties such as *Creator* to keep track of provenance. The *Tissue Block* is registered with the Biological Structure Ontology using annotation property *collides_with* with three PURL links to UBERON:0002015 (kidney capsule), UBERON:0001225 (cortex of kidney), and UBERON:0002189 (outer cortex of kidney) that all have GLB files with proper *Spatial Placement* in the 3D Reference Object Library.

Using annotation property *collides_with* (in lower left), we get PURL links to three anatomical structures—UBERON:0002015 (kidney capsule), UBERON:0001225 (cortex of kidney), and UBERON:0002189 (outer cortex of kidney)—which have corresponding GLB files that define their shape as well as their size, location, and rotation in relationship to the HRA 3D reference system. If anatomical structures have cell types and biomarkers listed in the ASCT+B table, it is possible to generate lists of cell types and biomarkers commonly found in these colliding anatomical structures. Note that the anatomical structure partonomy makes it possible to retrieve all sub-anatomical structures together with all the cells found in these.

##### Landmarks and extraction sites

There exist two types of spatial objects that ease tissue registration via the RUI (see **Fig. 1** and the Applications section). Landmarks such as the ‘bisection line’ for the kidney help align tissue blocks with real-world or man-made objects. Predefined tissue extraction sites used in GTEx can be assigned to many tissue blocks (e.g., in situations when tissue is always extracted from the very same spatial location in a human donor body). Both types of spatial objects are represented as a *Spatial Entity* that has been tagged as a member of an *Extraction Set*. Each *Spatial Entity* is associated with a 3D reference organ that has properties such as ‘organ owner sex’ and ‘left/right’ which uniquely identifies the organ. Examples of 43 *e*xtraction sites from four consortia are given in **Table 3**.

##### ASCT+B partonomy

The *‘part_of’ (BFO:0000050)* property is used to define the partonomy of anatomical structures and cell types. For example, in the kidney ASCT+B table (v1.2), we find *part_of* relationships between 1) anatomical structures:

> *cortex of kidney* ***SubClassOf*** *part_of* ***some*** *kidney*

*outer cortex of kidney* ***SubClassOf*** *part_of* ***some*** *cortex of kidney*

and 2) between a cell type and an anatomical structure, for example,

> *glomerular mesangial cell* ***SubClassOf*** *part_of* ***some*** *glomerular mesangium*

Note that the ASCT+B tables use *located_in* relationships between cells and anatomical structures to make the tables easier to author. In CCFO, *ccf_located_in* is the annotation property recording what the authors say. If this relationship validates when compared to existing ontologies, then *part_of* is added as the formalized relationship. Some partonomies are more than 10 levels deep. Note that *part_of* is transitive (i.e., it applies between successive members of a sequence but also between any two members taken in order; e.g., if A is *part_of* B, and B is *part_of* C, then A is *part_of* C) making ontology reasoning very efficient.

##### ASCT+B biomarker set

If cell types have biomarkers that *characterize* them listed in the ASCT+B tables, then the class *Sequence Collection (SO:0001260)* and the properties *has_characterizing_marker_set (RO:0015004)* and *has_marker_component* are used to construct the ontology statement in CCFO, for example,

> *glomerular mesangial cell* ***SubClassOf*** *has_characterizing_marker_set* ***some*** (
>
> *Sequence Collection* ***and***
>
> *(has marker component* ***some*** *periostin)* ***and***
>
> *(has marker component* ***some*** *PIEZO2)* ***and***
>
> *(has marker component* ***some*** *ROBO1)* ***and***
>
> *(has marker component* ***some*** *ITGA8)* ***and***
>
> *(has marker component* ***some*** *PDGFRB)* ***and***
>
> *(has marker component* ***some*** *SYNPO2))*

The interpretation of this statement is that the six cell-type markers *periostin* (HGNC:16953), *PIEZO2* (HGNC:26270), *ROBO1* (HGNC:10249), *ITGA8* (HGNC:6144), *PDGFRB* (HGNC:8804), and *SYNPO2* (HGNC:17732) together are sufficient to distinguish cells of type *glomerular mesangial cell* (CL:1000742) from others found in the same anatomical context (here, the anatomical structure *glomerular mesangium* [UBERON:0002320]). This assertion might be made based on human expertise or on experimental data. An example of the latter is the lung atlas work by Travaglini et al.^45^ that found that a minimum of 3-6 differentially expressed gene biomarkers is sufficient to uniquely identify 58 transcriptionally unique cell types in the lung for the spatial locations and demographic groups that were sampled. Note that tissue samples from different spatial locations (e.g., those not yet sampled at single-cell level) or donor populations (older donors or different ethnicities) might have different cell-type populations and characterizing biomarker sets.

#### CCFO Data Specifications

The CCF ontologies provide the vocabularies for data specifications that ease data exchange across consortia (e.g., see HRA reference organ description example in **Fig. 2** or HRA tissue block description in **Fig. 3**).

The data specification diagram of major CCFO classes and interlinkages is shown in **Fig. 4**. Some class pairs are linked through inverse relationships (e.g., a *Donor provides Sample*; *Sample comes_from Donor*). Most relationships are of single cardinality (indicated by a 1 on the arrow), whereas some are of multiple cardinality (indicated by an asterisk). Examples of multiple cardinality include: a donor can provide multiple tissue samples, a sample can be used to generate multiple datasets, a tissue block can be subdivided into multiple tissue sections.

**Figure 4.**
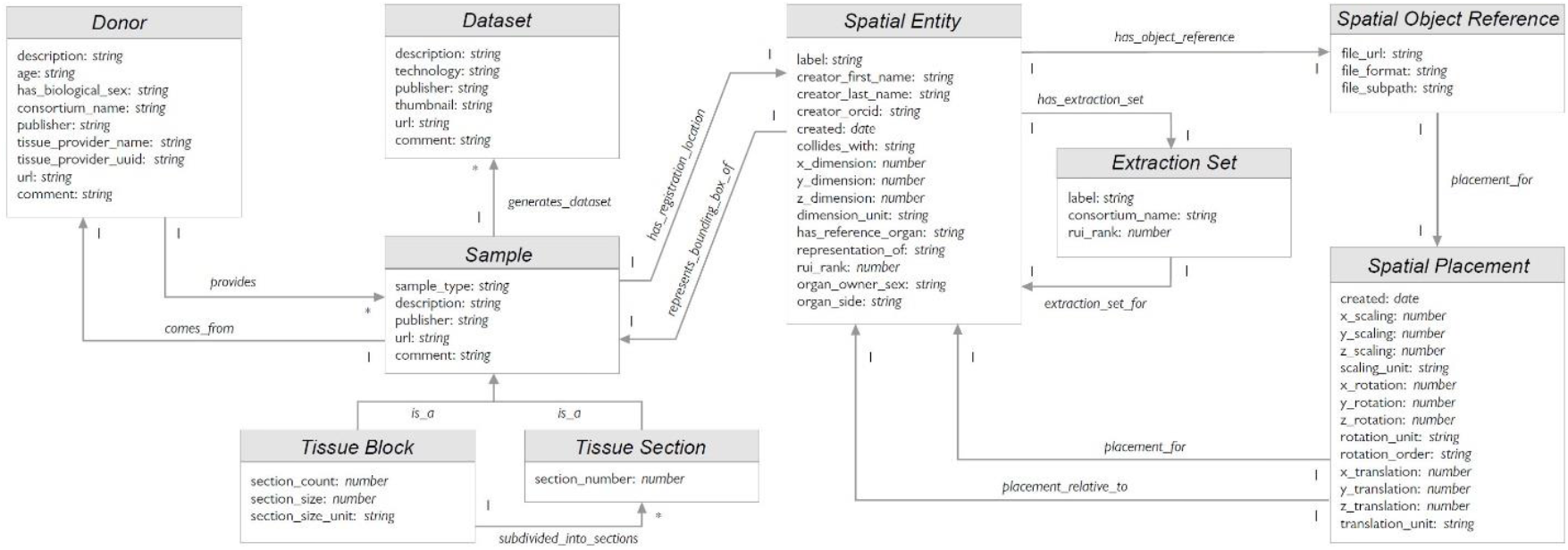
Data Specification Diagram. Major entities and their relationships are shown as used in JSON-LD. Note that a *Sample comes_from* a *Donor*; it is either a *Tissue Block* or a *Tissue Section*. A *Dataset* might be generated from a *Sample*. A *Sample* typically has a *Spatial Entity* which defines its size in relation to a *Spatial Object Reference* via a *Spatial Placement*.

The class *Spatial Entity* conceptualizes the notion of 3D objects in the CCFO. The coordinate framework for placing the 3D objects relative to each other is modeled through the class *Spatial Placement*. A 3D reference organ is a *Spatial Object Reference* with a *Spatial Placement* inside of the 3D HRA coordinate system (see example in **Fig. 2**). A *Tissue Block* is a *Sample* that *has_registration_location* defined via the *Spatial Entity* class relative to a *Spatial Object Reference* (see example in **Fig. 3**).

#### CCFO Editing, Building, and Release

A new release of the HRA is published every six months. The CCFO derived from the HRA data is published as a CCF.OWL graph database and associated CCF API shortly thereafter. Experimental datasets compliant with earlier versions of the HRA might need to be re-registered (e.g., if ASCT+B table terminology or 3D reference organs changed).

The “Standard Operating Procedure (SOP) Document for Ontology Editor”^46^ details how to (1) download and setup the development environment using Git, (2) add new reference organs to the CCFO, (3) update an existing reference organ in the CCFO, (4) build the CCFO, and (5) release the CCFO.

The general CCFO generation pipeline is shown in **Fig. 5**. The ASCT+B tables (in top left) are read by an external *Validation Tool*^47^ that compares whether anatomical structure and cell type terms are valid and if these relationships are true according to Uberon and CL/PCL using UberGraph^48^. All terms and relationships that already exist in Uberon and CL/PCL are added to the Biological Structure Ontology. When a term or relationship does not validate, the closest alternative match to another term in the ASCT+B Table is found and added. If no match is found, then ontology change request tickets (GitHub issues) are generated for missing anatomical structures, cell types, and relations for inclusion in existing ontologies. The result is a “partonomy” that reports *part_of* relationships between each pair of terms (anatomical structures to anatomical structures; cell types and anatomical structures (called *located_in* in the ASCT+B tables). A second pipeline uses the ROBOT tool^49^ to read the ASCT+B tables, augment them with information from reference ontologies (e.g., it adds metadata such as synonyms, definitions, or class hierarchies), and adds this additional data to the Biological Structure Ontology. Both processes generate weekly reports that are used to improve ASCT+B tables and to generate ontology change requests. Next, the *asctb2ccf*^50^ tool reads the validation tool report together with the original relationships from the ASCT+B table and generates a characterizing biomarker set for each cell type. *Spatial Object Reference* data (in middle left) is read by the *spatial2ccf*^51^ tool that generates instances of *Spatial Entity, Spatial Object Reference*, and *Spatial Placement* classes defined by the CCF Spatial Ontology. *Sample* registration data (in lower left) from HuBMAP, SPARC, KPMP, GTEx is available as LOD (see **Table 3** for examples). Each *Extraction Set* has a well-defined *Spatial Placement* (e.g., using the Registration User Interface in Applications section). Multiple tissue blocks can be associated with the same *Extraction Set* (e.g., GTEx has 29 *Extraction Sets* for more than 5,000 tissue blocks). The *asctb2ccf* tool reads the anatomical structure tags and generates Biological Structure Ontology data (e.g., predictions of cell types commonly found in these anatomical structures). The *spatial2ccf* tool reads spatial data and generates Spatial Ontology data. The *specimen2ccf*^52^ reads *Donor, Tissue Block*, and *Sample* data and generates Specimen Ontology data.

**Figure 5.**
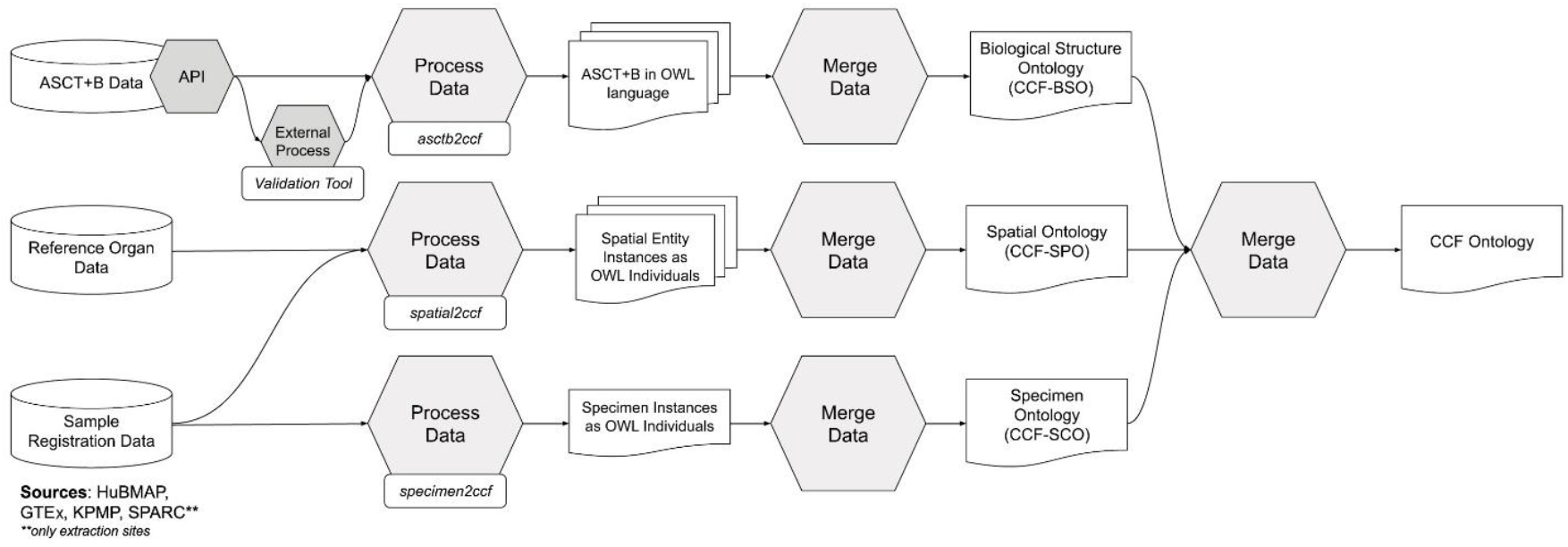
CCF Ontology Generation Pipeline. ASCT+B tables are validated and semantically enriched using the Validation Tool; in parallel, 3D Reference Organ data and crosswalk to ASCT+B tables are processed to compute the HRA. Sample Registration Data tissue blocks with RUI locations and anatomical structure tags can be retrieved for HRA construction or during HRA usage.

The CCF-Slim Ontology (ccf-slim.owl) captures HRA data exclusively (no experimental data), and it is used in the Registration User Interface (see Applications section).

## Data Records

The CCF.OWL v2.0.1 is available through the NCBI BioPortal Ontology Browser (https://bioportal.bioontology.org/ontologies/CCF) and the EBI OLS Ontology Browser (https://www.ebi.ac.uk/ols/ontologies/ccf). It is findable via https://purl.org/ccf/ccf.owl.

There are 266,576 triples in the graph database that holds the content of the CCF.OWL v2.0.1. The size of all 3D object GLB files is 2.56 GB.

CCF.OWL v2.0.1 represents major digital objects of the HRA v1.2, including 26 ASCT+B tables, 53 3D reference objects, and a crosswalk that associates anatomical structures in both. In addition, the data covers landmark organ sets which are used in the tissue registration user interface to facilitate tissue block placement in 3D organ models; these were not published with DOIs, but do appear in the CCFO.

CCFO classes and properties are listed in **Table 1**. Spatial entities representing 54 3D reference organs, 8 extraction sites, and 35 landmark extraction sites for a total of 97 are listed in **Table 2**. Experimental tissue data comprises 364 tissue blocks or registration sites and links to more than 5,000 tissue blocks and many more single-cell datasets, see **Table 3**.

**Table 2.**
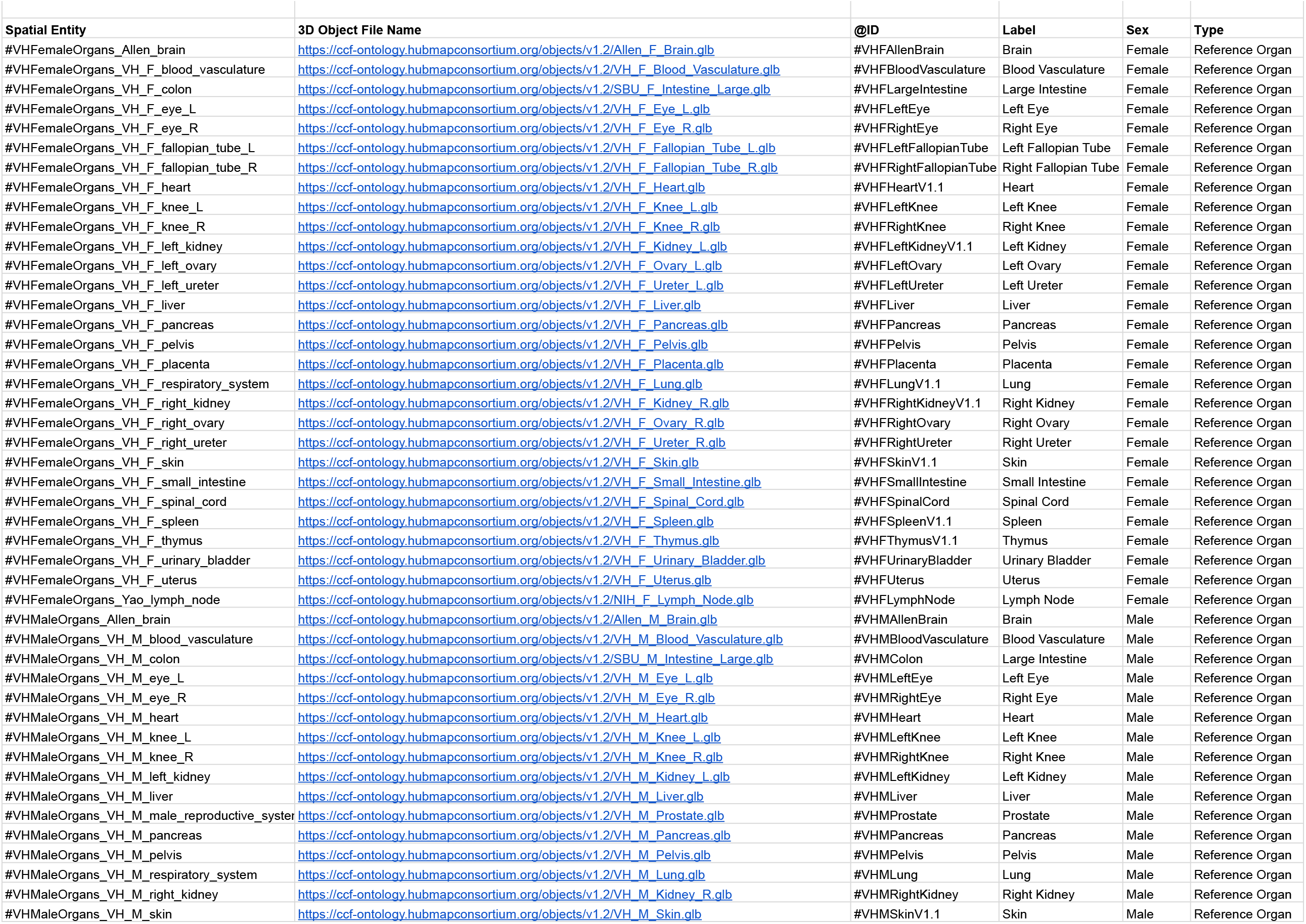
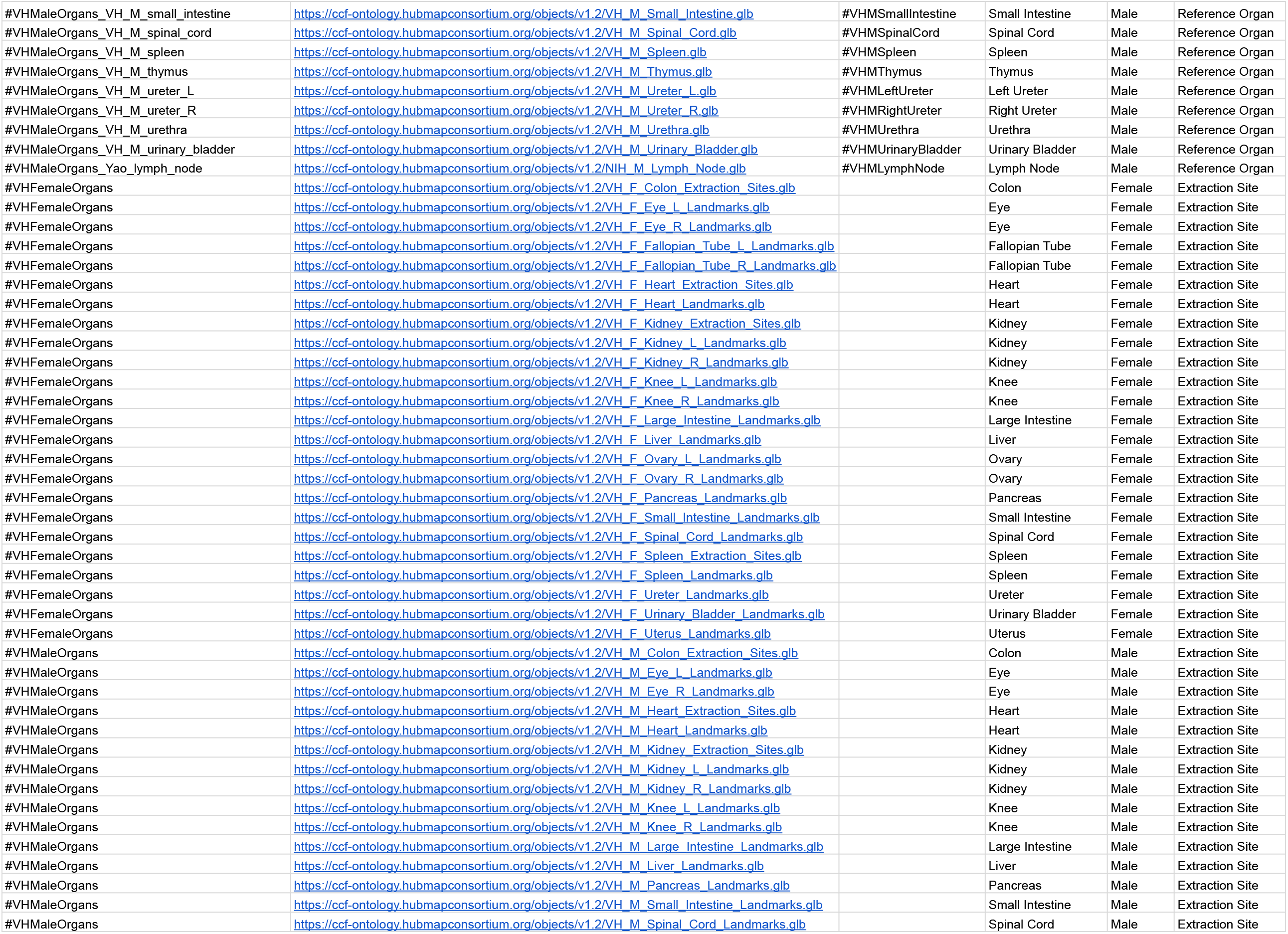
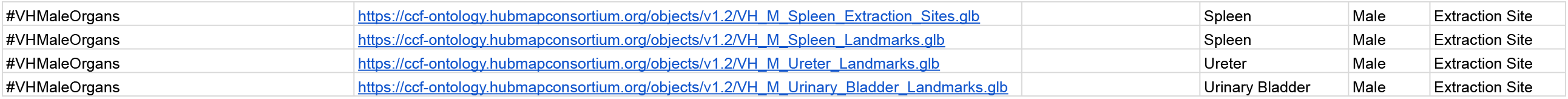
3D Reference Objects. The 3rd HRA release features 53 *Spatial Entities* of type *Spatial Object Reference* that are linked to anatomical structures in the ASCT+B tables plus 43 *Spatial Entities* of type *Extraction Site*. For each entity, we provide the spatial entity name, 3D object file name, ID and version number, organ label, sex, and type. Note that a landmark is a *Spatial Entity* that has been tagged as an *Extraction Set* for a specific *Extraction Site*, see **Table 3**.

**Table 3.**
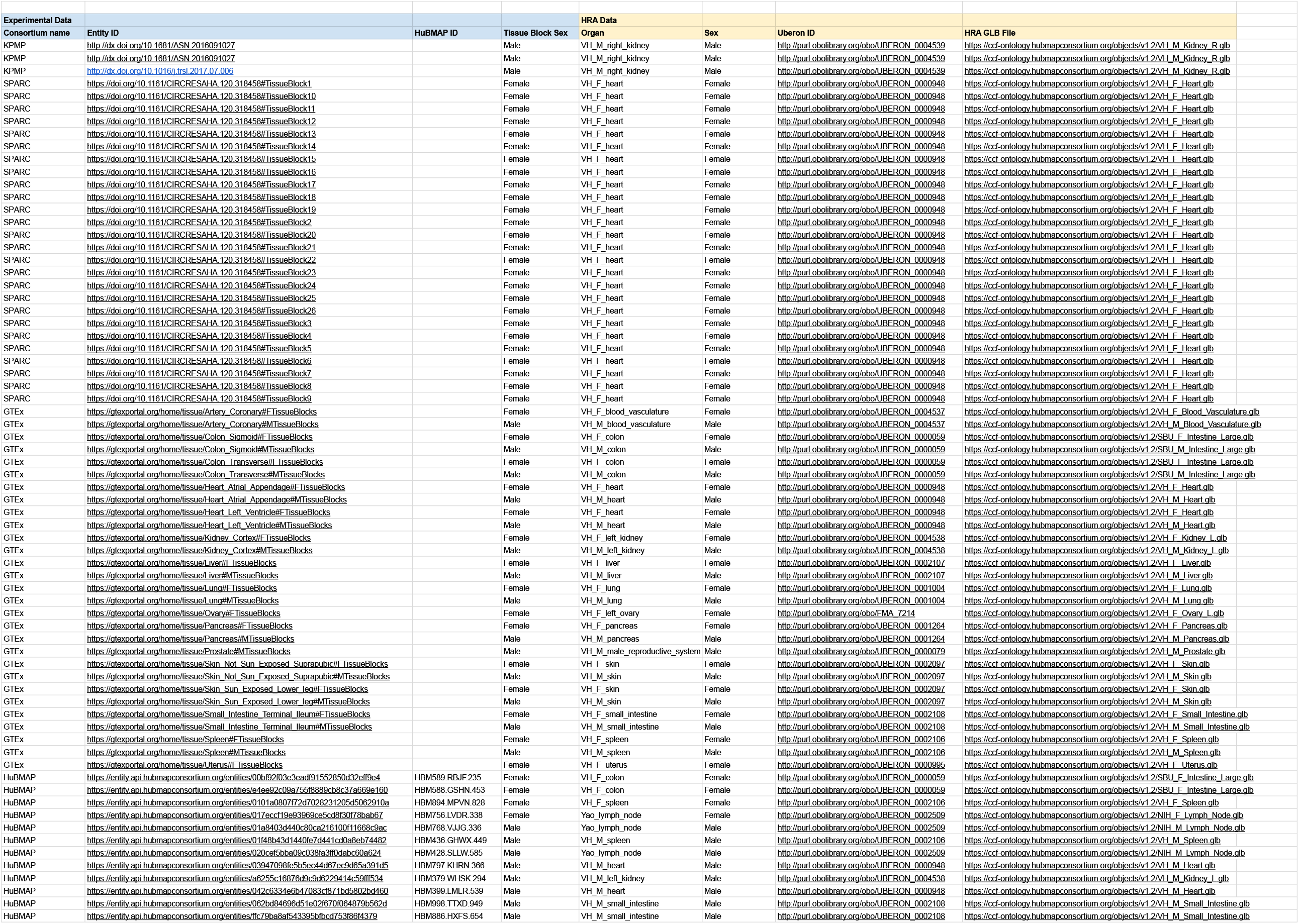
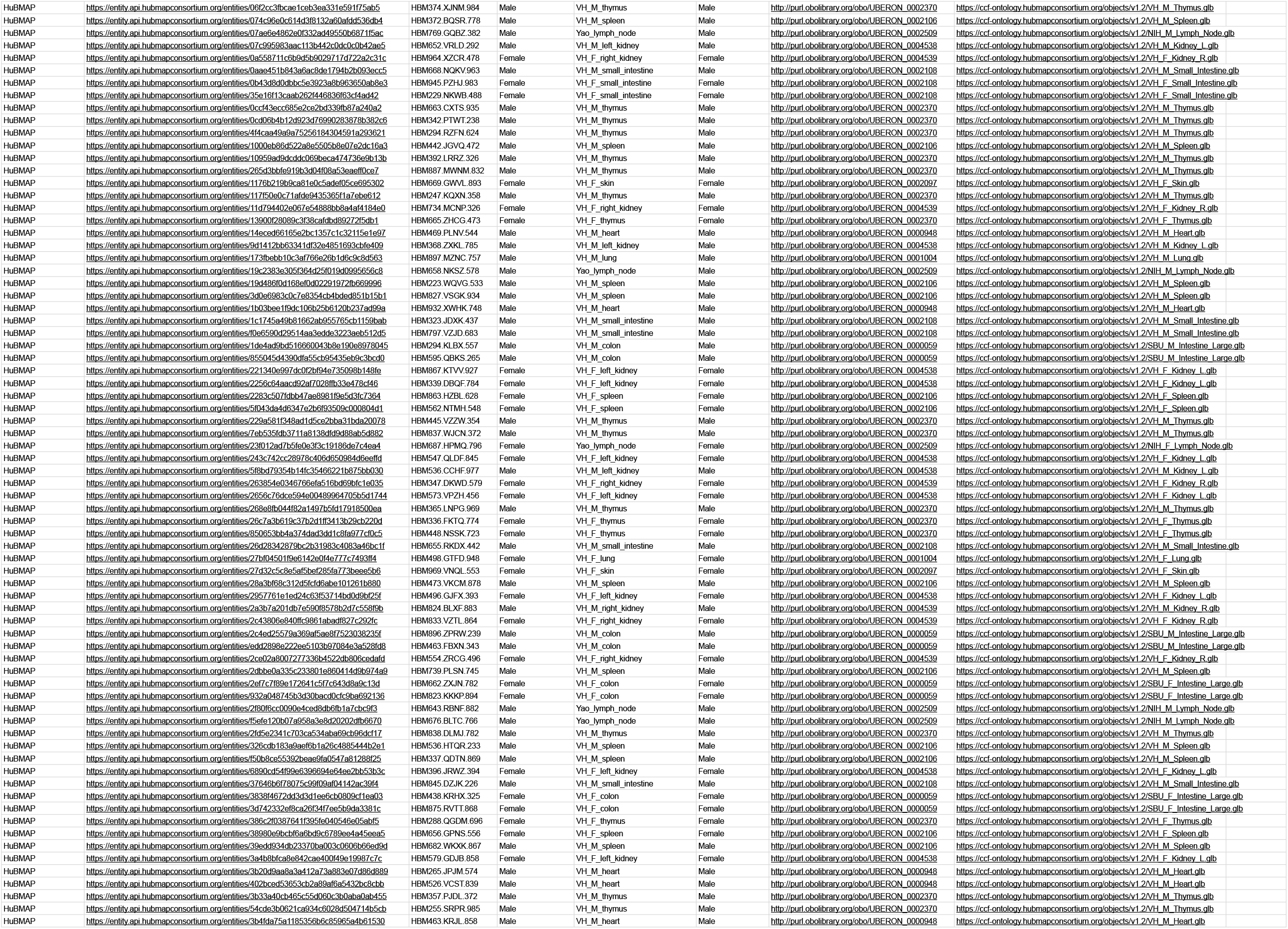
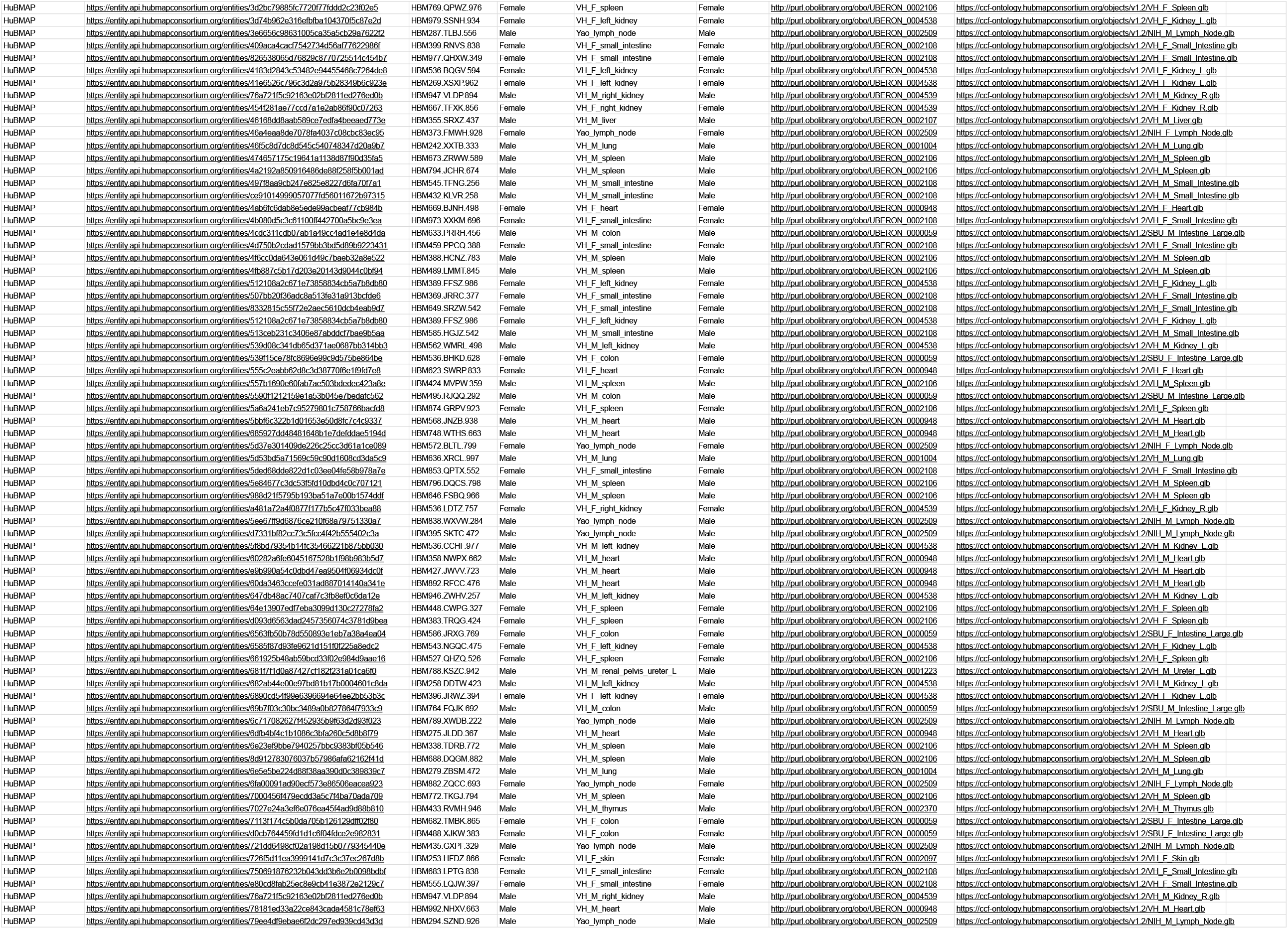
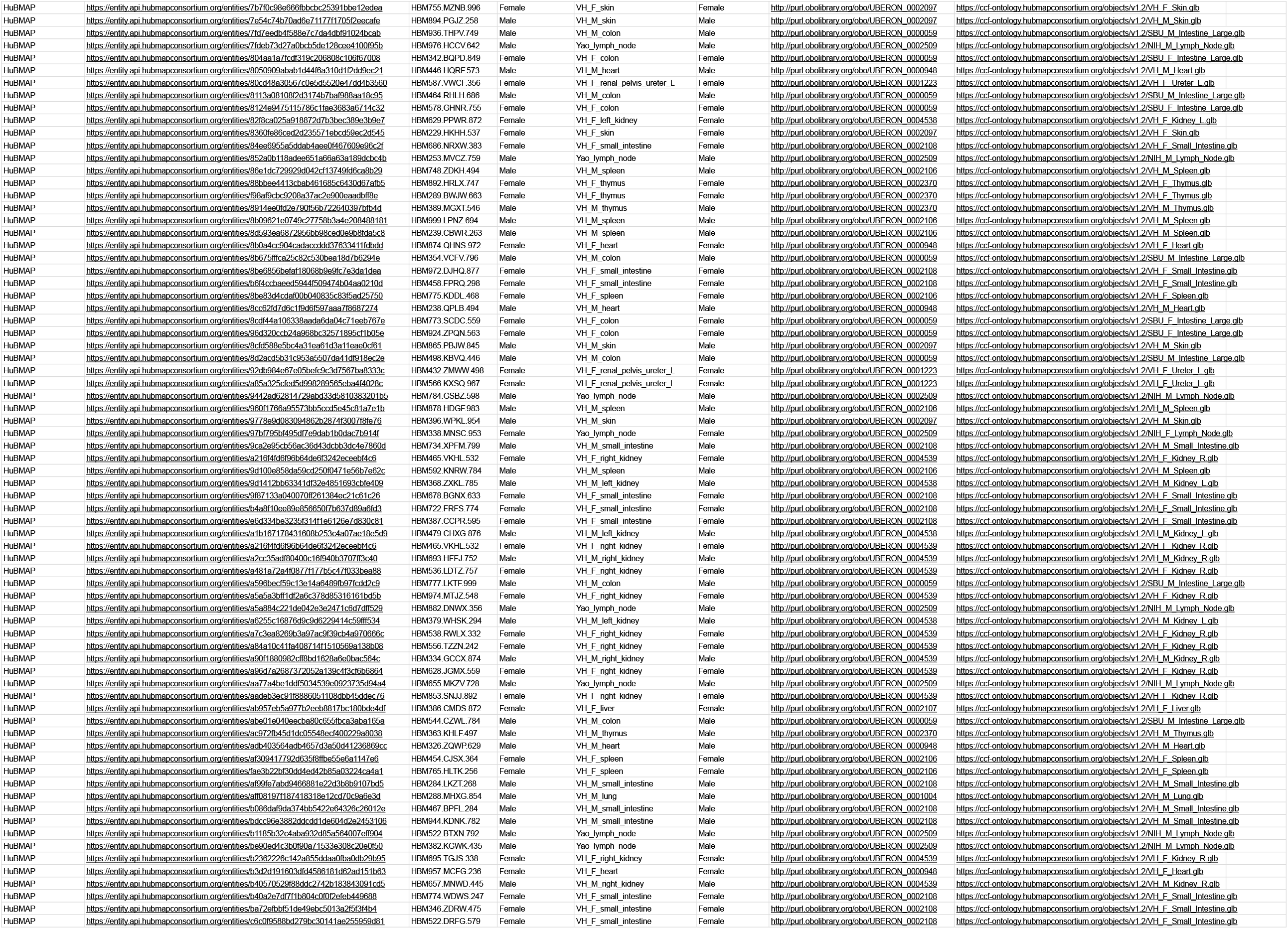
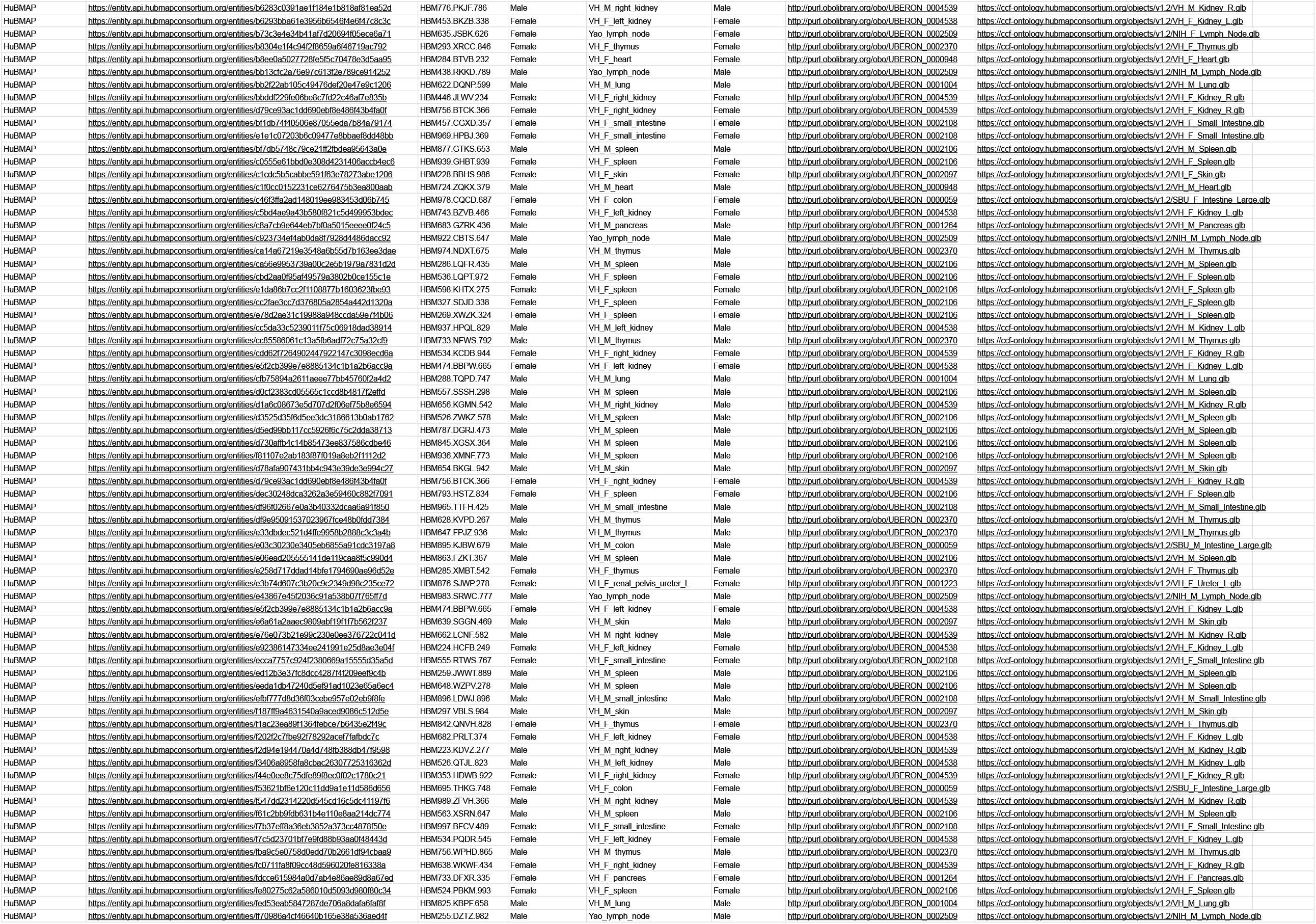
Registered Tissue Data. This table shows 364 *Extraction Site* entities that either represent the registration location for *Tissue Blocks* or *Registration Sites*. For each, we provide consortium name and entity ID, HuBMAP ID if available, and experimental data donor sex as well as HRA data such as organ name, sex, Uberon ID, and GLB file. The data comes from four consortia. HuBMAP data includes 306 tissue blocks, and many of these have derived tissue sections with one or more associated datasets. The 29 well-defined GTEx extraction set sites link to 5,397 datasets. The 26 SPARC blocks link to one paper while the three KPMP tissue blocks link to two data publications.

Weekly run term and relationship validation reports are available at https://github.com/hubmapconsortium/ccf-validation-tools/tree/master/reports.

Documentation and standard operating procedures have been published via the HRA Portal, https://hubmapconsortium.github.io/ccf.

## Technical Validation

### Source materials

The third HRA (v1.2) was authored by 114 unique experts that listed 185 publications with Digital Object Identifiers (DOIs). The 3D reference organs in this HRA were modeled using textbooks such as Gray’s Anatomy^53^, Netter’s Atlas of Human Anatomy^54^ and similar sources that are cited in the 3D model metadata. We hosted more than 50 organ-specific meetings with subject matter domain experts to discuss and review the evolving HRA. Experimental data generated by four consortia (HuBMAP, SenNet, KPMP, and GTEx) was mapped onto the HRA using 364 registered extraction sets that link to more than 5,000 tissue blocks.

### Comparison to Uberon, FMA, CL, HGNC

When comparing the ASCT+B tables on June 13, 2022 with Uberon (May 17, 2022 version) and Cell Ontology (CL, Feb 16, 2022 version), there were 1,620 terms that do not validate; 1,290 of these are from blood vasculature and peripheral nervous system. Many of the *part_of* relationships between anatomical structures validate, but few of the anatomical structures to cell type relationships validate (a large proportion is due to immune cells, which may or may not be tissue specific making it hard to model location in ontologies). For the extended blood vasculature tables, only 390 (41%) of the final 953 vessels in the HRA database are in Uberon. The FMA ontology has many terms that are missing in Uberon and we are in the process of adding those to Uberon. All 1,397 unique anatomical structures in the 26 organs 3D Reference Library link either to Uberon or FMA.

We use the CL as the main cell-type reference ontology. Recent technological advances in single-cell analysis make it possible to classify cell types based on their biomolecules in addition to their morphological attributes. This resulted in a large number of new cell types that are not yet recorded in CL. For example, there are 953 cell types covered by HRA (v1.2) that do not have a CL ID. We are using provisional cell-type terms and submitting new term request tickets so new terms can be added to CL.

As for biomarkers, the 3rd HRA features 2,842 total with 1,959 of type gene, 878 proteins, 3 proteoforms (in eye), 1 lipid (in placenta), and 1 metabolite (in eye). Of these, 447 biomarkers have provisional ontology IDs. Other gene and protein biomarkers are linked to HGNC.

### Applications

The CCF.OWL v2.0.1 and the CCF API are used by different applications. Four applications are detailed here and shown in **Fig. 6**. Each uses interlinked specimen, biological structure, and spatial data to provide unique functionality to users.

**Figure 6.**
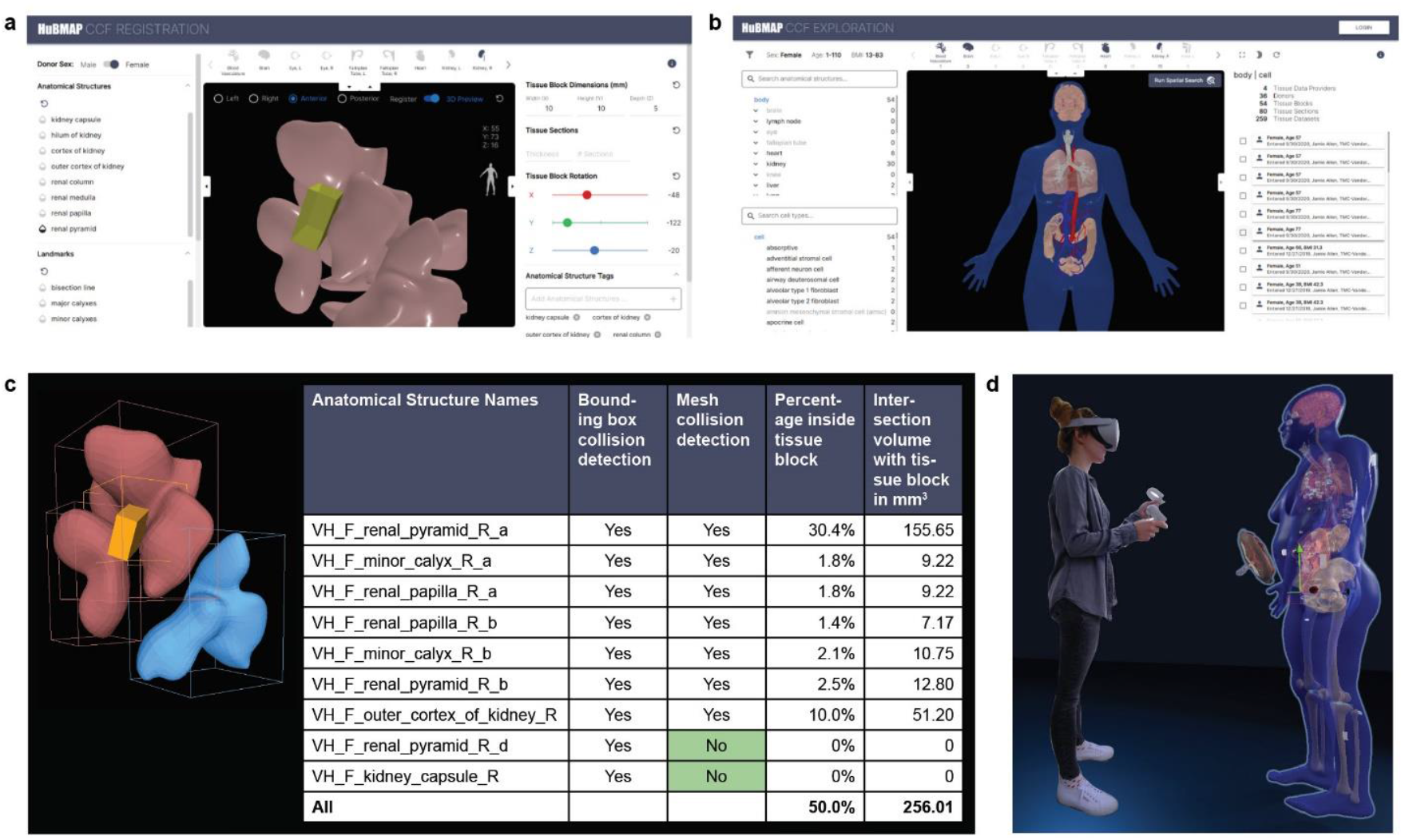
Applications. **a**. Registration User Interface (RUI) can be used to assign specimen, biological structure, and spatial metadata. **b**. Exploration User Interface (EUI) supports specimen, biological structure, and spatial search and exploration. **c**. Mesh-level collision detection improved semantic tagging. **d**. VR Organ Gallery with life-size 3D representations of the Visible Human Project dataset.

#### HuBMAP Portal

The Tissue Registration User Interface (RUI, see **Fig. 6a**) was integrated into the HuBMAP Ingest Portal to support the HRA-compliant registration of tissue samples^55^. RUI users first pick a 3D reference object that matches the organ from which tissue was extracted; then, they change the 3D size, position, and rotation of a virtual cube (in yellow) such that the values match the real-world tissue block properties—in relation to a 3D reference organ.

Collision detection (i.e., the intersection of the virtual cube with the reference organ anatomical structures is used to assign anatomical structure tags (see lower right of RUI). Spatial, biological structure, and specimen data is generated in JSON format. A stand-alone version of the RUI is available^13^ and data can be downloaded locally and shared freely as LOD so it becomes available via the CCF-API. The Exploration User Interface (EUI, see **Fig. 6b**) uses the CCF-API to let users explore RUI-annotated data using specimen, biological structure, and spatial attributes. For example, it is possible to filter tissue data collected by different consortia from different donors using properties such as donor sex and age, cell types, or spatial location. The code for both applications is freely available^11^. The EUI has been successfully integrated into the GTEx portal^26^.

#### Mesh-level collision detection

The current RUI performs collision detection at the bounding box level, generating more anatomical structure tags than desirable. Mesh-level collision detection checks if a tissue block collides with the 3D triangular meshes that define the surface of each reference anatomical structure. **Fig. 6c** shows an example of a tissue block (in yellow) colliding with three different renal pyramids in the female right kidney. Using bounding box collision, all three renal pyramids collide. Using mesh-level collision, only the two pink colored but not the blue colored renal pyramids collide. The data table shows Boolean values for collision detection using a traditional minimal bounding box (MBB)-based approach vs. mesh-based collision—differences are highlighted in green. The two columns on the right show the percentage and volume of different anatomical structures inside of the tissue block using mesh-level collision. Note that the sum total of AS volumes inside the tissue block is less than 100% as there is empty space between anatomical structures in the current 3D Reference Object Library.

To automate mesh-level collision detection, the HRA 3D Reference Objects were reviewed and re-meshed as needed to produce watertight meshes. For example, holes were filled using hole filling algorithms^56^ and non-manifold meshes were detected automatically but corrected manually by professional medical illustrators using 3D tools such as Zbrush^57^ and Maya^58^. Next, a multi-level spatial index^59^ was computed for each anatomical structure (i.e., the primitives such as faces, vertices and edges of each mesh) improving query performance by a factor of 10 compared with the traditional filter-refine paradigm used by PostGIS^60^. An in memory on-demand approach was used for data management, and an efficient collision detection algorithm^15^ was employed to compute the type and volume of anatomical structures within each tissue block. Last but not least, we computed the volume that is not inside any existing anatomical structures (e.g., that is outside of the organ, possibly indicating that a re-registration via the RUI is needed). All code was made available on GitHub^14^.

#### CCF Virtual Reality (VR) Organ Gallery

The VR Organ Gallery lets users explore the male and female human reference bodies in a 3D virtual environment using an Oculus Quest 2 virtual reality headset. The VR setup makes it possible to explore the HRA data across different scales—from encountering a whole body in real size (see **Fig. 6d**), zooming into single organs and studying their many anatomical structures, or examining cell-type populations within specific anatomical structures^12^. The Organ Gallery utilizes the CCF API to retrieve all 53 reference organs published in the third HRA release and 306 HuBMAP tissue blocks publicly available on August 25, 2022 (see **Table 3**). While the male and female reference body share many organs (such as brain, kidneys, colon), the male has the prostate, while the female body features the ovaries, fallopian tube, uterus, and placenta. Using HuBMAP IDs associated with each tissue block, additional specimen, provenance, and other data can be retrieved via the HuBMAP Entity API^61^ and made available via the application available on GitHub^62^.

## Limitations and future directions

The CCF.OWL v2.0.1 and the CCF API are used both in several production applications across consortia and in research and training. Yet, the current ontology model and API code have a number of limitations that have become obvious when using them in different contexts and for different use cases. Some modeling decisions (e.g., representing spatial entities and relationships using 3D meshes registered into a common coordinate system) have proven valuable while other decisions (e.g., representing landmarks as an *Extraction Site*) have been confusing for our team and other users and will be revised in future iterations. We discuss key limitations, lessons learned, and next steps here to empower other investigators who work on atlas design and/or applications that aim to facilitate access to large amounts of heterogeneous data.

### Coverage

CCF.OWL v2.0.1 does not cover all digital objects published in HRA v1.2. CCF.OWL v3.0 will aim to capture cell-type typologies, OMAPs, FTUs, and information on supporting scholarly publications and experimental data. We will model organs that have branching relationships with anatomical structures (e.g., blood vasculature, lymph vasculature, and peripheral nervous system) differently from those that have partonomic relationships (e.g., kidney or heart). We will add and expand the definitions of ontology classes and properties. In v3.0, all original entities and relationships from the ASCT+B tables will be preserved (as ccf property triples) as annotation property relationships, i.e., the can be queried separately from relationships derived from Uberon/CL or other reference ontologies.

### Improved validation

Currently, biological structure is validated against Uberon and CL. Going forward, we also validate biomarkers against HGNC and UniProt.References to entities that cannot be mapped will be added to issue trackers for eventual inclusion into existing ontologies.For example, since the publication of HRA v1.2, 127 terms from the provisional CL have been included and are now properly validating, and all 127 will be published in HRA v1.3. Note that the current set of 26 ASCT+B tables features links to LMHA for lung cell types and Provisional CL (PCL) for brain cell types; future revisions of the HRA will aim to replace them by links to CL.

### Data provenance

Novel experimental data can substantially change the HRA reference and mapping procedure. Hence, it is of utmost importance to capture the provenance of all data and code used in atlas creation and to recompile the HRA when new evidence becomes available. Going forward, we expect to see multiple biomarker sets (e.g., generated using different cell-type annotation tools) per cell-type population; these can be encoded using Boolean AND between all required biomarkers from one set and OR between biomarker sets; this will likely lead to ontology reasoner scaling issues. Future CCF.OWL versions will support complete provenance chains for all digital objects and code using W3C Provenance^63,64^.

### Spatial accuracy

To understand the limitations of tissue registration using the Registration User Interface (RUI), we performed a user study with 42 subjects^55^. Results show that RUI users can perform tissue block matching tasks at an average of 5.88 degrees rotation accuracy and 1.32 mm position accuracy (at an average of 22.6 seconds per task after 8.3 tasks in a series of 30 identical tasks). Going forward, we will implement Stage 2 registration using biomolecular data that is only now becoming available for some anatomical structures. Over time, with sufficient high-quality data from thousands of human donors, the HRA will become more stable and data registration more accurate.

### Proper separation between ontologies and data instances

CCF.OWL v3.0 will model HRA- relevant vocabularies, application ontologies, and references using the LinkML^65^ general purpose modeling language. The HRA will use CCFO to detail all specimen-related, structural biology, and spatial data. Experimental data from various consortia will be modeled using LinkML and advertised as 5-star Linked Open Data^66^. That is, the CCFO defines the vocabularies shared by HRA and experimental data; LinkML is used for the data specification. Digital objects (also called data instances) will not be part of the CCFO but will be made available via a separate data management system.

### Novel use cases

A series of qualitatively new use cases will be developed to showcase the value of combining HRA with other LOD data—e.g, from the Gene-Centric Common Fund Data Ecosystem (CFDE) Knowledge Graph^67^, the STRING - Protein-Protein Interaction Networks Functional Enrichment Analysis database^68^, and the Scalable Precision Medicine Oriented Knowledge Engine (SPOKE) graph database that federates more than 30 open datasets into a public data commons of health relevant knowledge^16,69^.

### Scaling up

Efficient data management, spatial queries and algorithm implementations are needed to scale up to the 37 trillion cells that make up the human body^2^ and to truly capture human diversity. CCF data is a core component that will be accessed highly frequently by many database queries. To guarantee fast, real-time query response of CCF data, we will continue to monitor and improve the implementation through techniques such as in-memory data management, hybrid relational and graph-based storage, and parallel processing.

### Documentation and training

This paper details the CCFO and the CCF API for the first time. Going forward, we plan to publish simple examples of how the CCFO is currently used (or will be used) in different applications. We will organize hands-on workshops that train other experts to run proper SPARQL queries against the CCF.OWL graph database or use the standard HTTP CCF-API to support a whole ecosystem of collaborative and compatible APIs, libraries, and user interfaces.

## Usage Notes

The CCF API supports programmatic access to CCF.OWL v2.0.1 data; for exemplary queries see the companion website at https://cns-iu.github.io/ccf.owl-paper-supporting-information.

The CCF.OWL v2.0.1 data and API are used in the HuBMAP and GTEx portals, the Registration and the Exploration User Interfaces, 3D mesh-level collision detection, and the VR Organ Gallery.

We welcome others to join our effort in constructing a user-friendly and useful HRA. Please sign up for the ASCT+B Working Group at https://iu.co1.qualtrics.com/jfe/form/SV_bpaBhIr8XfdiNRH to receive updates and meeting invites.

## Code Availability

All data and standard operating procedures are released under Creative Commons Attribution 4.0 International (CC BY 4.0). All code was released under the MIT License and can be accessed in GitHub at https://github.com/hubmapconsortium/ccf-ontology.

### ASCT+B APIs

API Endpoint (includes interactive documentation): https://asctb-api.herokuapp.com

API Documentation: https://hubmapconsortium.github.io/ccf-asct-reporter/docs/api

OpenAPI specification: https://asctb-api.herokuapp.com/asctb-api-spec.yaml

### CCF-API

API Endpoint (includes interactive documentation): https://ccf-api.hubmapconsortium.org

API Documentation and OpenAPI specification: https://ccf-api.hubmapconsortium.org

API Database backend is N3.js: https://github.com/rdfjs/N3.js

Code to instantiate/use CCF Database: https://github.com/hubmapconsortium/ccf-ui/tree/main/projects/ccf-database

### CCF-User Interfaces

https://github.com/hubmapconsortium/ccf-ui

### Validation Tools

https://github.com/hubmapconsortium/ccf-validation-tools

## Acknowledgments

We thank the more than 200 authors that contributed to the ASCT+B table authoring and review process; Kristen Browne and Heidi Schlehlein for creating the 3D reference objects; Devin M. Wright and the HuBMAP team for registering 306 tissue blocks and Peter Hanna from SPARC, Seth Winfree from KPMP, and Kristin Ardlie from GTEx for close collaborations on using the CCF in their consortia efforts. We deeply appreciate work by Tracey Theriault for figure design, Todd Theriault for copy editing, and Medina Sydykanova and Naval Pandey for procuring images. This research has been funded by the National Institutes of Health Human BioMolecular Atlas Program (HuBMAP) award OT2OD026671, Cellular Senescence Network (SenNet) Consortium Organization and Data Coordinating Center (CODCC) award 1U24CA268108-01, National Institute of Diabetes and Digestive and Kidney Diseases (NIDDK) Kidney Precision Medicine Project grant U2CDK114886, NIH U01CA242936, and Common Fund Data Ecosystem (CFDE) OT2 OD030545.

## Author contributions

BWH, JH, EMQ, DOS, MAM, KB designed the CCF ontologies; BHW, JH, DOS implemented the ontology generation pipeline; JH, MAM validated the specimen ontology; EMQ, DOS, ARC validated the biological structure ontology; BWH, AB, CH, FW used the CCF ontology in different applications; BWH, JH, EMQ, AB, LC, FW, DOS, MAM, KB wrote the paper.

